# The first nested association mapping (NAM) population for outbreeding Italian ryegrass reveals candidate genes for seed shattering and related traits

**DOI:** 10.1101/2025.10.22.683898

**Authors:** Jenny Kiesbauer, Christoph Grieder, Meril Sindelar, Linda Helene Schlatter, Daniel Ariza Suarez, Steven Yates, Ingrid Stoffel-Studer, Dario Copetti, Bruno Studer, Roland Kölliker

## Abstract

Nested association mapping (NAM) populations are a powerful tool for investigating the genetic control of agronomically important traits and have been successfully used in many inbreeding crops. Here, we present the first NAM population established in an outcrossing forage grass species, Italian ryegrass (*Lolium multiflorum* Lam.), to dissect the genetic control of seed shattering. The NAM population was based on 23 diverse and one common founder plants and consisted of 708 F_2_ individuals. Reduced-representation sequencing (ddRAD) of the 708 F_2_ individuals, combined with whole genome sequencing data of the 24 founder plants, yielded a total of 3,199,253 SNPs that were used for population structure analysis, parentship analysis and genome-wide association studies. Phenotypic data for seed shattering and seed yield-related traits, collected in three year x location environments, showed high phenotypic variance within the NAM population. A total of seven QTL were identified for seed shattering, seed yield, spike length, flag leaf length and flowering time. Within these QTL regions, one candidate gene for seed shattering and three candidate genes for flowering time were identified. For seed shattering, the significant SNP association within the gene *chr7.26897,* known to be in involved in ripening-related pathways, explained 10.03% of the phenotypic variance. These candidate genes identified provide valuable targets for functional validation and demonstrate the effectiveness of NAM populations for elucidating the genetic architecture of complex traits in outcrossing forage grasses.

## Introduction

More than 50% of the European agricultural land area is covered by permanent and temporary grasslands (FAO, 2020). Italian ryegrass (*Lolium multiflorum* Lam.) is one of the most abundant grasses used for forage production in temperate regions. It is a naturally diploid (2*n*=2*x*=14), allogamous species with an estimated genome size of 2.6 Gb (Kopecký et al., 2010). Due to its self-incompatibility, both natural and breeding populations show high levels of heterozygosity which makes genomics-assisted breeding strategies challenging (Byrne et al., 2015; Frei et al., 2021; Knorst et al., 2019). Efficient breeding strategies require a comprehensive understanding of the genetic control of the target traits. In Italian and perennial (*L. perenne* L.) ryegrass, quantitative trait mapping using biparental mapping populations has been widely used as a strategy to elucidate the genetic control of traits such as disease resistance (Jo et al., 2008; Schejbel et al., 2007; Studer et al., 2007), heading date, spike length, spikelets per spike (Byrne et al., 2009), fatty acid composition (Hegarty et al., 2013), lodging resistance (Inoue et al., 2004), winter survival (Paina et al., 2016) or seed yield and fertility traits (Studer et al., 2008). However, biparental populations have several limitations. First, the low number of recombination events occurring during population development may lead to a low mapping resolution (Scott et al., 2020). While increasing the number of individuals can partially mitigate this issue, this approach is constrained by time, costs and availability of resources. Second, the allelic diversity represented by only two parental plants may not sufficiently capture the breadth of natural variation, thereby limiting the number of detectable quantitative trait loci (QTL) and their transferability to other populations (Yu et al., 2008). Multi-parent populations are particularly powerful to investigate the genetic architecture of complex traits. Nested association mapping (NAM) populations combine the advantage of a high allelic diversity and a high statistical power to detect the alleles present in the population (Yu et al., 2008). In general, NAM populations are developed by crossing multiple diverse founder plants with a single common founder plant, followed by self-pollination of the progenies resulting in a set of recombinant inbred lines (RILs). To successfully establish NAM populations, several prerequisites must be fulfilled. First, the founder plants should be as genetically diverse as possible to maximize allelic variation for the traits of interest. Second, the higher the number of diverse founder plants included in the NAM population, the higher the statistical power to detect a QTL will be (Stich, 2009). Third, each family within the NAM population should consist of a sufficient number of individuals to provide adequate representation and ensure robust statistical analyses. In major crops, NAM populations have been successfully used to identify genomic regions associated with highly complex traits, each having small effects (Gangurde et al., 2020; Herzig et al., 2019; Poland et al., 2011; Yu et al., 2008). To date, NAM populations in major crops were mainly designed for inbreeding species based on the development of RILs from F_1_ individuals (Gage et al., 2020; Gangurde et al., 2020; Herzig et al., 2019; Poland et al., 2011; Yu et al., 2008) or from backcrossed-derived RILs and doubled haploids (Kitony, 2022). In outcrossing species like Italian ryegrass, RILs cannot be produced via self-pollination. However, a modified version of a NAM population in Italian ryegrass at the F_2_ level may be produced by crossing one common founder plant with many diverse founder plants, followed by open-pollination among the F_1_ individuals.

Intensive breeding efforts for Italian ryegrass only started in the 1980s with a focus on agronomic performance such as persistence, forage quality and, most importantly, biomass yield (Jung et al., 1996; Marshall & Wilkins, 2003). When a newly developed cultivar is ready for commercialization, the main bottleneck for forage grass cultivars at present is seed multiplication (Casler & Vogel, 2020; Falcinelli, 1999). Seed yield depends, among other factors, on morphological traits of the flowering head such as spike length or flag leaf size (Abel et al., 2017; Elgersma, 1990; Schierenbeck et al., 2024). Potential seed yield defines the maximum seed yield achievable under optimal conditions and depends mainly on the number of ovules per area present at flowering (Falcinelli, 1999). However, up to 40% of the potential seed yield can be lost due to unsuccessful pollination, fertilization, seed setting and seed development as well as by the shattering of seeds (Falcinelli, 1999). Unsuccessful pollination and fertilization can occur by non-optimal pollen supply during flowering (Boelt & Studer, 2010). A successful fertilization is only possible if at least one of the alleles at the *S* and *Z* loci from the two-loci gametophytic self-incompatibility system are different between the male (pollen) and female (ovule) determinant (Cornish et al., 1979; Hayman, 1956; Lundqvist, 1954; Rohner et al., 2023). After successful fertilization, seed set is defined by embryo and endosperm growth, which is highly sensitive to weather conditions (Elgersma & Śnieżko, 1988). In ryegrasses, shortly before or during harvest, a substantial number of seeds can be lost due to seed shattering (Kiesbauer et al., 2023; Maity et al., 2021). Also, variation in flowering time, which is often high in Italian ryegrass breeding populations, can lead to inhomogeneous ripening of the seeds and can, therefore, potentially increase seed shattering. Thus, seed shattering is an important trait in ryegrasses, resulting in cultivars with low seed yield. In wild plant species, seed shattering is advantageous to disperse seeds from the maternal plants, resulting in a higher species fitness (Li & Gill, 2006; Maity et al., 2021). However, in grain crops such as rice (*Oryza sativa* L.) and sorghum (*Sorghum bicolor* L. Moench), seed shattering was one of the key traits selected for during domestication (Fuller, 2007; Harlan, 1975), but was never a selection target in Italian ryegrass, which are primarily bred for biomass yield. To date, more than 60 QTL associated with seed shattering have been published for rice (Wu et al., 2023). Therefore, seed shattering appears to be a quantitative trait controlled by multiple genes with different effect sizes. Due to the advantages already described, such as displaying high allelic diversity and a low population structure, a NAM population is an ideal tool to elucidate the genetic basis of such a highly quantitative trait as seed shattering. The objectives of this study were therefore a) to develop and genetically characterize the first F_2_ NAM population for Italian ryegrass, b) to phenotypically evaluate the NAM population for seed shattering and other seed yield-related traits and c) to identify genomic regions linked to seed shattering and related traits as a basis for genomic-assisted selection.

## Materials and methods

### Establishment of the nested association mapping population

The common founder plant (MC), which is characterized by relatively high seed shattering, was selected from the Swiss cultivar ‘Rabiosa’ and has been used to produce a high-quality phased reference genome (Chen et al., 2025). The 23 diverse founder (MD1–MD23) plants were selected from cultivars, ecotypes, and breeding material (Table 1). Data on seed shattering was available from previous experiments for MD1-MD5, but not for MD6-MD23.

**Table 1.** List of the common (MC) and diverse founder plants (MD1-23) used for the establishment of the F2 nested association mapping population for Italian ryegrass *(Lolium multiflorum* Lam.). Name indicates the accession from which the individual founder plants were selected from. Pollen donor indicates how many times an F1 plant originally generated from the biparental cross between the common and the respective diverse founder plant was identified as the pollen donor for the 708 F2 plants using a correlation approach.

| M | Name | Population type | Seed shattering phenotype | Pollen donor |
| --- | --- | --- | --- | --- |
| C | Rabiosa | Cultivar | High seed shattering | - |
| D1 | Sikem | Cultivar | Low seed shattering | 19 |
| D2 | Shakira | Cultivar | Low seed shattering | 37 |
| D3 | Srl1_LI1115_3501-02 | Breeding material | High seed shattering | 32 |
| D4 | Srl2_LI1266_3413-10 | Breeding material | High seed shattering | 15 |
| D5 | Srl3_LI1245_3426-10 | Breeding material | High seed shattering | 17 |
| D6 | LI9901 | Breeding material | Unknown | 21 |
| D7 | LI9902 | Breeding material | Unknown | 24 |
| D8 | Arth_Chraebel | Ecotype | Unknown | 43 |
| D9 | Cadenazzo | Ecotype | Unknown | 40 |
| D10 | Castel_S.Pietro | Ecotype | Unknown | 36 |
| D11 | Egg | Ecotype | Unknown | 20 |
| D12 | Tremona | Ecotype | Unknown | 27 |
| D13 | Davinci | Cultivar | Unknown | 24 |
| D14 | Majesty | Cultivar | Unknown | 29 |
| D15 | Oryx | Cultivar | Unknown | 38 |
| D16 | Rangifer | Cultivar | Unknown | 48 |
| D17 | LI0085 | Breeding material | Unknown | 36 |
| D18 | LI0505 | Breeding material | Unknown | 35 |
| D19 | LI0615 | Breeding material | Unknown | 31 |
| D20 | LI0905 | Breeding material | Unknown | 13 |
| D21 | LI0915 | Breeding material | Unknown | 34 |
| D22 | LI1015 | Breeding material | Unknown | 20 |
| D23 | LI1135 | Breeding material | Unknown | 22 |

In a first step, all the 23 MD were crossed to MC, resulting in 23 F_1_ families. Seeds were germinated for two days in petri dishes, and seedlings were established in the greenhouse for approximately four weeks. To confirm the F_1_ offspring, the plants were verified using simple sequence repeat markers specific to each of the parental founders, and the confirmed plants were transplanted to the field. In the subsequent year, four to five tillers from 18 randomly chosen F_1_ plants from each of the 23 F_1_ families were cut shortly before flowering for a randomized open pollination in the greenhouse. Seeds were harvested separately from each of the 414 F_1_ plants. The F_2_ seeds were then germinated for two days in petri dishes, and seedlings were established in the greenhouse for approximately four weeks. Two seedlings from each of the 414 F₁ parents were randomly selected, resulting in an initial NAM population of 828 F₂ plants. After transplantation to the field, 120 plants did not survive. Consequently, the NAM population used in this study consisted of 708 F₂ plants (Figure 1).

**Figure 1.**
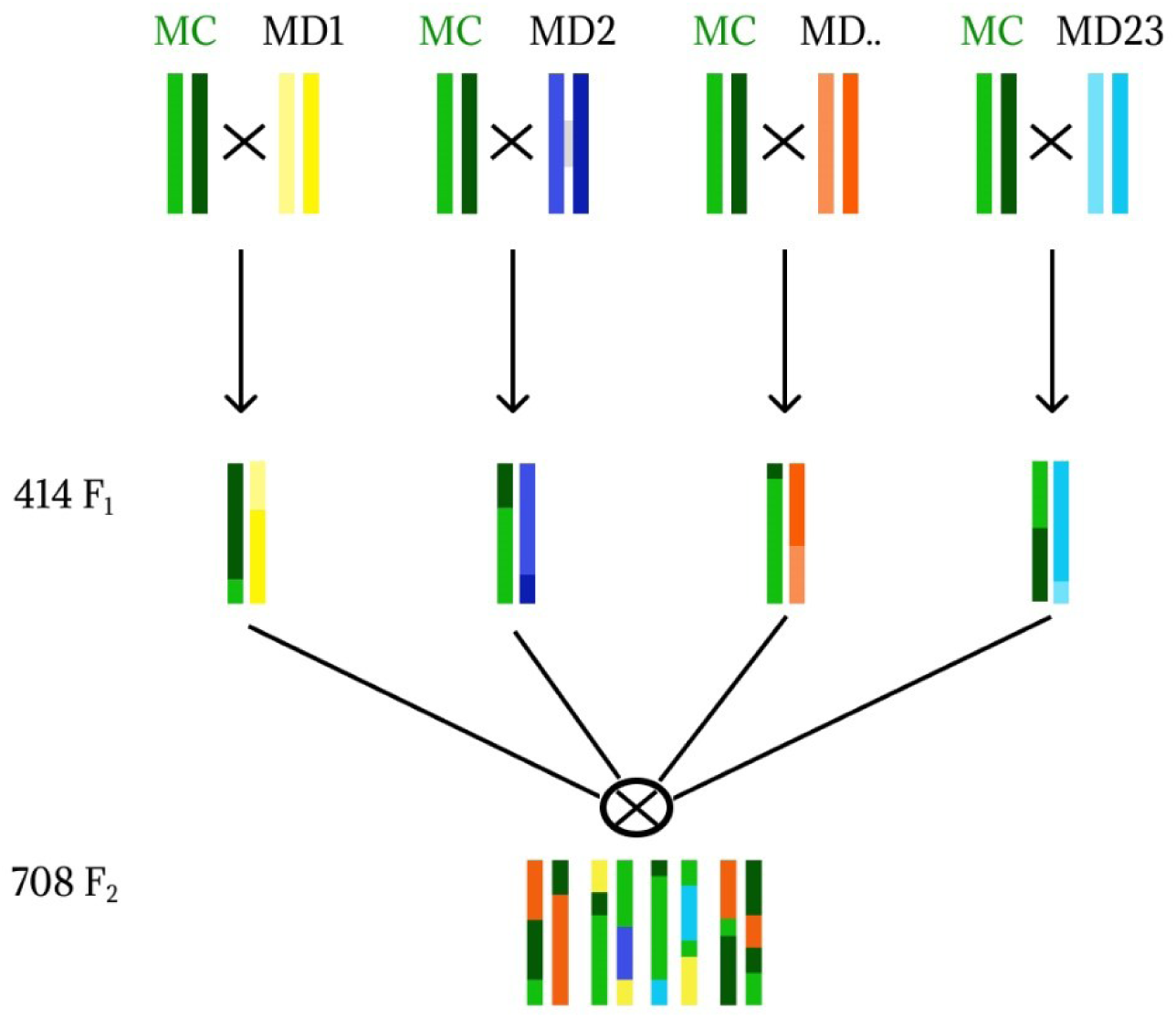
Development of the nested association mapping (NAM) population for Italian ryegrass (Lolium multiflorum Lam.). Biparental crosses of 23 diverse (MD) with one common founder (MC) plant resulted in a F1 population of 414 plants. From these 414 F1 plants, two seeds per plant were sown and used for open pollination, resulting in 828 F2 plants. In total, 708 F2 plants survived and constituted the final NAM population

### Genotyping and SNP calling

For each of the 708 F_2_ plants and the 24 founder plants, a segment of 2 cm of mature plant leaf was sampled and stored at -80°C for DNA extraction. After grinding the samples with a TissueLyserII (Qiagen, Hilden, Germany) at 25Hz for 40 seconds using two grinding beads (Tungsten Carbide Beads, 3 mm, Qiagen, Hilden, Germany), DNA was extracted with the Mag-Bind® Plant DNA DS 96 Kit (Omega Bio-tek, Inc. Norcross, GA, USA) according to the recommended protocol by the manufacturer. DNA quality was checked on a 1.2% agarose gel (220 V, 300 mA, 20min). DNA quantity was measured with Quant-iT™PicoGreen™ ds DNA Assay Kit (ThermoFisher Scientific, Waltham, Massachusetts).

Whole-genome sequencing (WGS) was performed for the 24 founder plants, using 150 bp paired-end sequencing on the Illumina NovaSeq 6000 platform at the Functional Genomics Center Zurich, Zurich, Switzerland, aiming for a minimum raw sequencing depth of 15× coverage per genotype. In addition, double digest restriction site associated DNA (ddRAD) sequencing libraries using the restriction enzymes *EcoRI* and *TaqI* (New England Biolabs Ipswich, MA, USA) were prepared from the extracted DNA for all 24 founders plants and the 708 F_2_ plants following the protocol of Peterson et al. (2012). The ddRAD libraries were sequenced with 150 bp paired-end reads using the Illumina NovaSeq 6000 platform by Novogene (Cambridge, United Kingdom). The reads were demultiplex using Stacks (v2.65; Catchen et al., 2013), allowing up to one mismatch in the adapter sequence. Adapter tails from ddRAD and WGS data were clipped with HTStream (v1.3.0; https://github.com/s4hts/HTStream). The reads from WGS and ddRAD were mapped to the reference genome of the MC (Chen et al., 2025) using Bowtie2 (v2.5.1; Langmead & Salzberg, 2012). The mapped WGS and ddRAD reads were used for SNP calling using NGSEP (Tello et al., 2023).

A genotypic matrix derived from the WGS data of the 24 founder plants was filtered for genotype calls with a quality score above 30, minimum read depth (DP) of three and no missing data per SNP. This matrix was phased using Beagle (v5.4; Browning et al., 2018), using a biparental genetic linkage map for *L. multiflorum* (Chen et al., 2025) and the options ‘burnin=9’ and ‘iterations=36’. A second genotypic matrix derived from ddRAD data contained the 24 founders and the 708 NAM F_2_. This matrix was obtained by genotyping the WGS-derived list of SNPs with the option ‘-knownVariants’ of NGSEPs *SingleSampleVariantsDetector* module. This matrix was similarly filtered for genotype qualities above 30, minimum DP of three, and at least 1% of the accessions genotyped per SNP.

The missing data in the ddRAD matrix was imputed with Beagle, providing the WGS-derived matrix as the reference panel, the same biparental genetic linkage map (Chen et al., 2025) and the options ‘window = 200’ and ‘imp-states = 250’ following the recommendations of (Pook et al., 2020) for low-diversity datasets. Finally, the imputed matrix was filtered to remove SNPs with minor allele frequencies below 0.02. The predicted effect of the SNPs on the gene models of the reference genomes was annotated with snpEff (v5.0e; Cingolani et al., 2012).

The imputed ddRAD data was used for principal component analysis (PCA) using TASSEL (v 5.0; Bradbury et al., 2007), to calculate pairwise measures of linkage disequilibrium (LD) using a sliding windows approach of 50 markers per chromosome and to conduct GWAS. The R package LDcorSV (v1.3.3; Mangin et al., 2012) was used to correct the LD measures for kinship relationships in the population (*r^2^_v_*). The LD decay was visualized by estimating the pairwise regression values of *r^2^_v_* on the physical distance of the markers using loess regression (R function ‘loess’), which smoothens numerical vectors with a span value of 0.75 (R Core Team, 2024).

### Parentage analysis to identify pollen donors

Parentage analysis of the F_2_ plants was performed using two different methods. First, the relationship between two individuals was calculated using KING (Manichaikul et al., 2010) and the imputed genotypic matrix consisting of 3,199,253 SNPs. KING robustly calculates pairwise kinship coefficient, without being influenced by sample composition or population structure. The second method was based on Pearson’s correlation of the genotypic data between individual F_2_ plants and the 23 MD genotype, hereafter referred to as correlation analysis. This analysis used the genotypic data of each SNP coded as 0, 1, 2 for homozygous reference, heterozygous and homozygous alternative alleles, respectively.

### Field trials and phenotyping

For the first trial (ZH20), two clonal replicates of each F_2_ plant were transplanted to the field in Zurich (47.427 °N, 8.516 °E) in autumn 2020. In spring 2021, plants were cut for the first time after completion of heading. During the second growth, the plants were fixed to bamboo sticks and phenotyped for seed shattering and seed yield-related traits as described below. After phenotyping and harvesting in autumn 2021, the 708 plants were dug out and divided into four clonal replicates, which were then planted in two additional trials in Zurich (47.427 °N, 8.516 °E; ZH21) and in Oensingen (47.284 °N, 7.730 °E; OE21), with two clonal replicates per trial. These were then phenotyped during the second growth of the following year (2022). In all three trials, an alpha design with two replicates and incomplete blocks of ten plants being planted in a row was used (Piepho et al., 2006). Planting distance was 40 cm within, and 50 cm between rows of 10 plants. Management of trials and assessment of traits were the same in all three environments. Each trial was fertilized in spring once with 50 kg ha^- 1^ nitrogen and with 50 kg ha^-1^ nitrogen after each cut. Traits were assessed during the second growth, where time of heading and flowering time (measured in days after April 1) were also determined for each plant. As soon as a plant started flowering, length of the flag leaf and length of the inflorescence were assessed as the average of the measurements from the four highest tillers. Once all plants flowered, all inflorescences from each plant were bagged into perforated plastic bags (Sealed Air, Cryovac, Charlotte, North Carolina, USA, 330 mm × 750 mm) and plants were tied to bamboo sticks. The harvesting date of plants was adjusted by flowering time and temperature sum (cumulated average day temperature as measured 5 cm above ground from a nearby weather station). In each trial, the harvest time of the earliest flowering plant was determined based on seed maturity, and the temperature sum accumulated since its start of flowering was noted. All other plants were harvested on the day when the temperature sum accumulated since their start of flowering reached the same value as for the earliest flowering plant. Corresponding temperature sums were 618 °C, 598 °C and 555 °C for the trials ZH20, ZH21 and OE21, respectively. At harvesting, the bagged inflorescences were cut with a sickle, carefully shaken three times, and the inflorescence with the remaining seed was placed in a second bag. Before the plants were placed in the second bag, the occurrence of stem rust was assessed on each whole plant previously placed in a bag rated on a scale from 1 to 9, defined as: 1 = no rust disease, 2 = trace of rust, 3 = 5%, 4 = 10%, 5 = 25%, 6 = 40%, 7 = 60%, 8 = 75% and 9 = more than 75% of the plant is covered with rust (Schubiger & Boller, 2016). Both bags were put in a drying cabinet under constant air flow at 23 °C for three days. Empty seeds, tillers as well as debris were removed from the viable seeds using an airflow-separator (Saugluft-Stufensichter T2, Baumann, Saatzuchtbedarf Waldenburg Germany). Using a digital scale (New Classic MS, Mettler Toledo, Columbus, Ohio, USA, accuracy 0.01 g), the weight of shattered and non-shattered seeds was measured. We defined seed shattering (%) as:

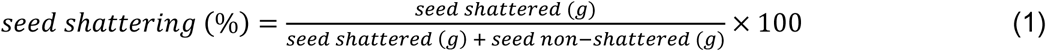

The total seed yield was calculated as the sum of seed shattered (g) and seed non-shattered (g).

### Statistical analysis of phenotypic data

To analyze the phenotypic data across environments, the following linear mixed model was used,

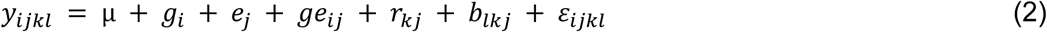

where *y_ijkl_* represents the measurement for trait *y* on a single plant basis, *µ* denotes the overall mean, *g_i_* the effect of genotype *i*, *e_j_* the effect of environment *j*, *ge_ij_* the interaction effect of genotype *i* with environment *j*, *r_kj_* the effect of the complete block (replicate) *k* nested within environment *j*, *b_lkj_* the effect of incomplete block *l* nested within complete block *k* and *ε_ijkl_* the residual error. First, all factors except *b_lkj_* were treated as fixed to estimate best linear unbiased estimators (BLUEs) for each genotype across all environments. Second, the factors *g_i_* and *ge_ij_* were additionally taken as random to estimate genotypic and genotype-by-environment interaction variance. To adjust for any effect of stem rust infection on seed yield, stem rust scoring was included as an additional covariate (fixed effect) in model (2). For estimation of the respective variance components, the stem rust scoring was taken as a random effect. The broad sense heritability (*h^2^*) was calculated as

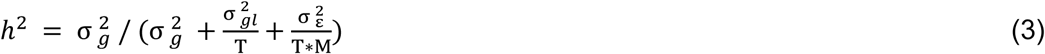

where 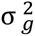, 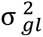 and 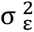 are the genotypic, genotype-by-environment interaction and residual variance, respectively. T = 3 is the number of environments and M = 2 is the number of replicates per environment.

All phenotypic data analyses were conducted using RStudio v2025.02.28 and R v4.4.3 (R Core Team, 2024). Package dplyr (v1.1.4; Wickham et al., 2023) was used for data manipulation, lme4 (v1.1.35.5; Bates et al., 2015) for fitting mixed-effect models, lmerTest (v3.1.3; Kuznetsova et al., 2017) to display significance of factors anova and emmeans (v1.10.4; Lenth, 2023) for calculating the BLUEs.

### Genome-wide association analysis

Genome-wide association mapping was conducted with a total of 3,199,253 biallelic SNPs, and GAPIT (v3; Wang & Zhang, 2021), using a mixed linear model (MLM; Yu et al., 2006), a multiple locus mixed linear model (MLMM; Segura et al., 2012), BLINK (Huang et al., 2019) and FarmCPU (Liu et al., 2016). For each model, the covariables included the first three principal components derived from the genotypic matrix consisting of 3,199,253 SNPs. Manhattan and QQ-plots were generated from GAPIT results using the R packages qqman (v0.1.9; Turner, 2018). Significant associations were declared when the *p*-value for each SNP marker was smaller than the Bonferroni-corrected threshold, which was calculated with a genome-wide type I error rate of 0.05 or a threshold of *p* = 1.0 × 10^-5^. The four different methods used for association analysis (i.e., MLM, MLMM, BLINK and FarmCPU) revealed partly inconsistent results regarding significantly associated loci. Only marker trait associations that were significant with the BLINK or/and the FarmCPU model and that showed a clear peak (however not always significant) in the Manhattan plots using the MLM and MLMM models were further investigated.

### Identification of candidate genes

LD analysis was used to determine LD decay, with *r^2^*= 0.1. A confidence interval of 14.4 kb around SNPs significantly associated with traits was applied for candidate gene identification in the reference genome of the MC (Chen et al., 2025). To find information on the putative function of these candidate genes, the gene sequence from the reference genome was compared to other crop genomes (*L. rigidum, L. perenne, Oryza sativa*) using BLAST+ (Camacho et al., 2009) in NCBI using the nucleotide database (Sayers et al., 2021).

## Results

### Population structure, parentage analysis and LD of the NAM population

In total, the imputed genotypic matrix of the NAM population consisted of 3,199,253 SNP. The number of SNPs per chromosome ranged from 349,793 (chromosome 5) to 619,781 (chromosome 4). As the Italian ryegrass genome, more precisely the genome of MC, has an estimated size of 2.46 Gb (Chen et al., 2025), the genome-wide average marker distance was 768.93 bp. The first two axes of a PCA of the entire genotype matrix explained 1.72% and 1.50% of the total genetic variation, respectively, and revealed no obvious population structure or clustering based on the founder plants (Figure 2). One small cluster was identified, which contained four plants potentially originating from foreign pollination.

**Figure 2.**
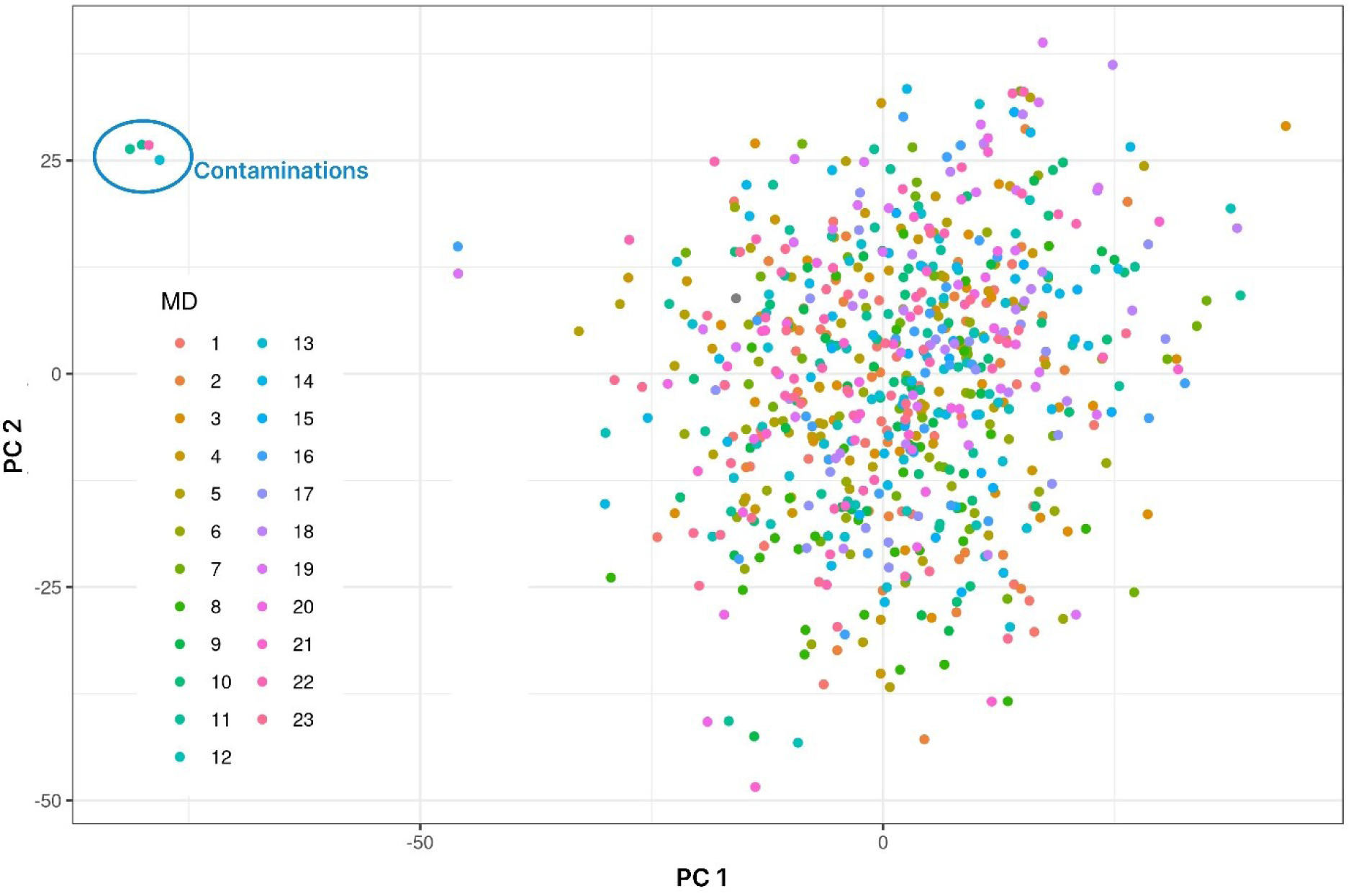
Genetic structure of the nested association mapping population consisting of 708 F2 plants. The first principal component (PC1) explained 1.72%, the second (PC2) 1.50% of the total genetic variation within the population. The maternal origin (i.e., the F1 family and the respective founder genotype) is indicated by different colors (diverse founders MD1–MD23). The four plants showing large deviation from the others, most likely being a result of foreign pollen during open pollination among F1 plants, were labelled as contaminations and excluded from all further analyses.

Due to the design of the NAM population, each F_2_ plant traces back to maximally three different founder plants, of which one is always the MC. One of the two MD is known from the pedigree information (i.e., the MD that was the parent of the F_1_ plant from which F_2_ the seed was harvested; the mother plant) and the other corresponds to the MD of the unknown pollen donor F1 plant. Correlation analysis and parentage analysis using KING identified four plants as outliers (Figure 3) with unknown founder plants, corresponding to the four plants in the small cluster identified by PCA (Figure2; on the left). Hence, these four genotypes were removed for subsequent GWAS. Using KING, the two maternal founders (MC and MD) could be correctly verified in 90.62% of the tested cases. In 5.20% of the cases, KING indicated different MD as maternal founders than those defined by the pedigree. Of the remaining 4.18% of cases tested, only one MD could be clearly assigned as maternal founder. In particular, the KING method failed to assign the founder MD11. In total, 7 out of 12 instances KING was not able to identify MD11 as MD. As the correlation analysis proved more accurate, correctly assigning 96.10% of the tested cases, the correlation method was used instead of KING to assign maternal and paternal MDs (Table 1, Figure 3). However, due to their complementary approaches, both approaches provided a robust strategy for identifying the parents of each individual in the NAM population. If the pollinations were equally distributed among the MDs, each of the 23 MDs would contribute on average to 30.8 progeny. Within the population, the average of pollination per founder was 28.7. From the 23 MDs, MD20 and MD4 contributed the least (with 13 and 15 times, respectively) and MD8, MD9 and MD16 contributed the most (with 43, 40 and 48 times, respectively), to the pollination of the F_1_ plants.

**Figure 3.**
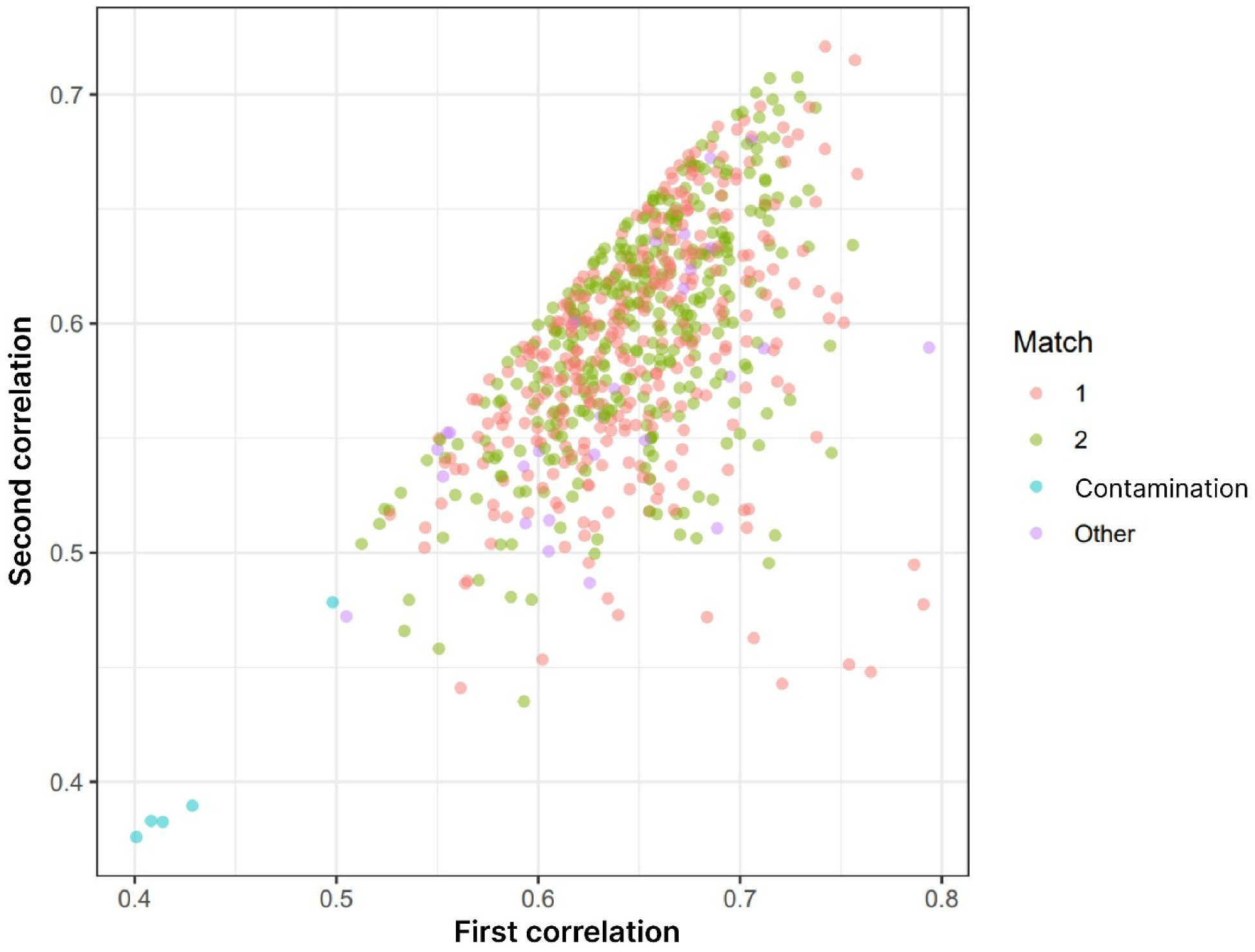
Biplot showing the highest (*x*-axis) and second highest (*y*-axis) correlation coefficient of every F2 plant to one of the 23 diverse founder plants. Match indicates whether the highest (1, red dots) or second highest (2, green dots) correlation was the maternal founder known from pedigree information. Contaminated (blue dots) means that the correlation between the founders and the tested F2 NAM plants showed a very low correlation. Other (purple dots) represent F2 plants that were correlated to one of the founder plants, but not to those derived from pedigree information.

The LD decay with physical distance between SNPs showed slightly lower LD decay on chromosome 7. Across all seven chromosomes, on average a LD decay within 14.4 kb has been detected with a threshold of *r^2^* = 0.1 (Figure 4).

**Figure 4.**
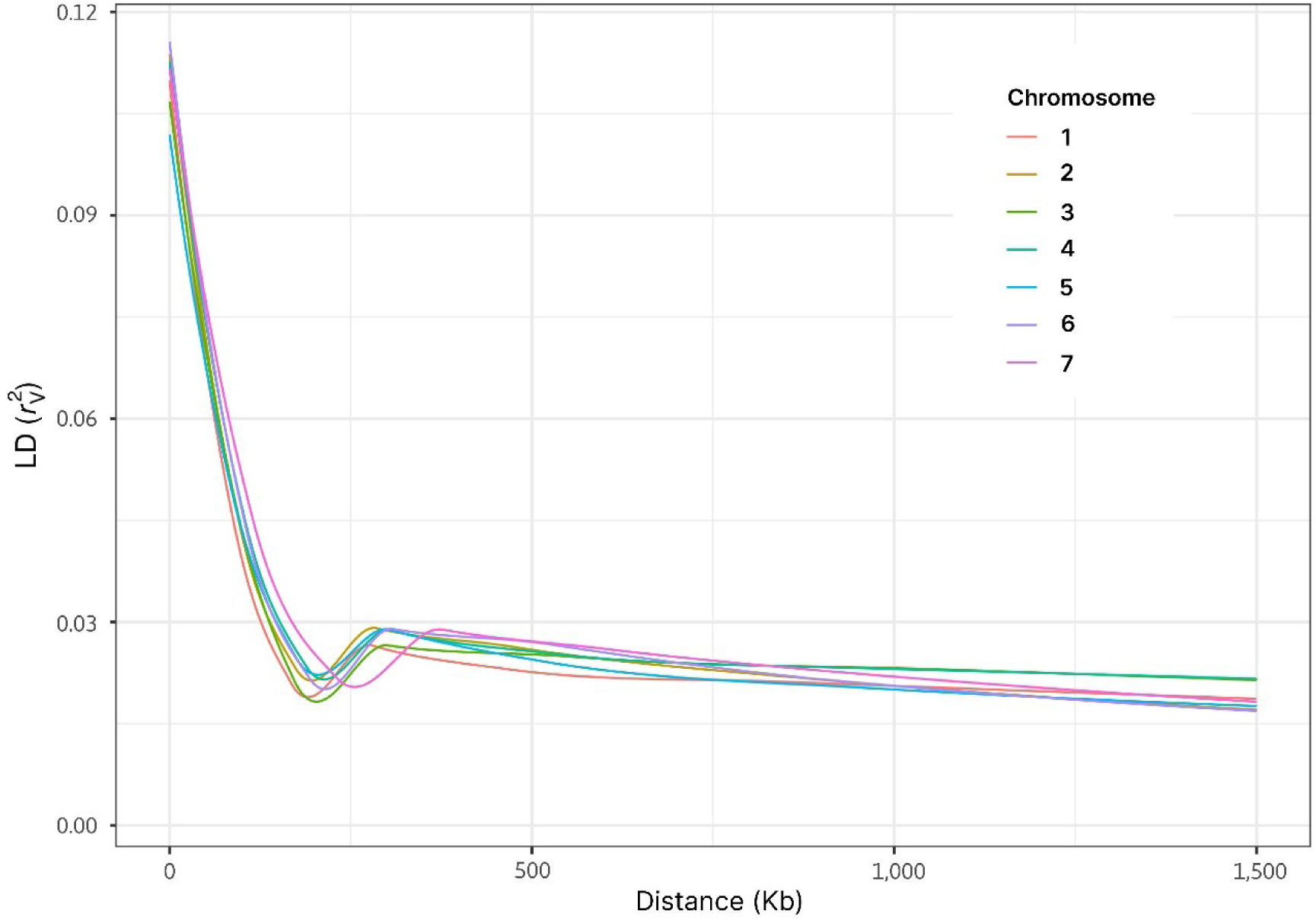
Genome-wide linkage disequilibrium (LD) decay estimated per chromosome. The LD (*r*^2^v) measured between pairs of polymorphic markers is plotted against the physical distance (bp) between these loci. The pairwise measures of LD were calculated using a sliding window of 50 markers. The colored lines display LOESS regressions for each chromosome.

### Phenotypic variation for seed shattering and seed yield-related traits

On average, the NAM F_2_ plants started flowering after 83.61 days (measured in days after first of April), had a flag leaf length of 16.97 cm, a spike length of 21.26 cm, 18.49 spikelets per spike and shattered seeds at a rate of 37.09% with a total seed yield of 11.69 g. The highest standard deviation was seen for seed shattering (σ = 14.99), followed by seed yield (σ = 4.52). In general, high to moderate heritabilities were observed for the phenotypic traits, ranging from *h^2^*= 0.92 for spike length to *h^2^* = 0.68 for flowering time (Table 2). For spike length, seed shattering per plant and seed yield per plant, the largest part of the variance was explained by the genotype, while residual variance was largest for the traits flowering time and flag leaf length. The variance explained by the genotype × environment interaction was very small for spike length (9% of genotypic variance) but larger for the other traits (29-59% of genotypic variance; Table 2). Correlations among the phenotypic traits were only weak to moderate (Table 3), showing highest phenotypic correlation coefficients (*r_p_*) between spike length and flag leaf length (*r_p_* = 0.62). Seed yield showed a moderate correlation to flag leaf length (*r_p_* = 0.41). Seed shattering only showed weak correlations with other traits, like flowering time (*r_p_* = -0.13), flag leaf length (*r_p_* = 0.14) or spike length (*r_p_* = 0.08).

**Table 2.** Mean (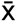) and standard deviation (σ) of the best linear unbiased estimators per genotype, heritability (*h^2^*), genotypic variance component (σg^2^), genotype × environmental interaction variance component (σ_gl_^2^) and residual variance component (σ_ε_^2^) for the traits seed shattering per plant (%), seed yield per plant (g), time of flowering counted in days after 1st of April, spike length (cm), number of spikelets per spike and flag leaf length (cm) are displayed. For σ_gl_^2^ and σ_ε_^2^ the relative values compared to σg^2^ are given in brackets.

| Trait | $\bar{x}$ | $\sigma$ | $h^2$ | $\sigma_g^2$ | $\sigma_{gi}^2$ | $\sigma_e^2$ |
| --- | --- | --- | --- | --- | --- | --- |
| Seed shattering per plant (%) | 37.09 | 14.99 | 0.84 | 194.29 | 56.73 (0.29) | 121.44 (0.63) |
| Seed yield per plant (g) | 11.69 | 4.52 | 0.79 | 15.04 | 4.92 (0.33) | 14.04 (0.93) |
| Flowering time (days) | 83.61 | 1.66 | 0.68 | 1.74 | 1.02 (0.59) | 2.80 (1.61) |
| Spike length (cm) | 21.26 | 3.14 | 0.92 | 3.61 | 0.46 (0.13) | 1.64 (0.45) |
| Number of spikelets per spike | 18.49 | 2.00 | 0.90 | 8.98 | 0.80 (0.09) | 2.86 (0.32) |
| Flag leaf length (cm) | 16.97 | 2.28 | 0.79 | 4.05 | 1.16 (0.29) | 4.13 (1.02) |

**Table 3.** Pearson’s correlation coefficients (*rp*) based on best linear unbiased estimators of genotypic means among the traits seed shattering per plant (%), seed yield per plant (g), flowering time (days), spike length (cm), number of spikelets per spike and flag leaf length (cm). Significance levels are shown as * = *p*-value < 0.05, ** = *p*-value < 0.01 and *** = *p*-value < 0.001.

| Trait | Seed yield per plant (g) | Flowering time (days) | Spike length (cm) | Number of spikelets per spike | Flag leaf length (cm) |
| --- | --- | --- | --- | --- | --- |
| Seed shattering per plant (%) | 0.11** | -0.13*** | 0.08* | 0.01 | 0.14*** |
| Seed yield per plant (g) |  | -0.29*** | 0.34*** | 0.20*** | 0.41*** |
| Flowering time (days) |  |  | -0.19*** | -0.04 | -0.21*** |
| Spike length (cm) |  |  |  | 0.50*** | 0.62*** |
| Number of spikelets per spike |  |  |  |  | 0.26*** |

### Genome-wide association mapping

With our NAM population design, we were able to identify seven QTL on chromosomes 1, 4, 6 and 7 associated with flowering time, spike length, flag leaf length, seed shattering and seed yield per plant (Figures 5-10). In these QTL regions, two candidate genes were identified for seed yield, three for seed shattering, two for flag leaf length and six for flowering time. All SNPs significantly (*p* < 0.005) associated with the corresponding traits found with BLINK or/and FarmCPU and consistently detected a clear peak in the Manhattan plots with the other models (MLM and MLMM) are listed in Table 4 and shown in Supplementary Figure S5. For each trait, genes identified within 14.40 kb (up- and downstream) regions of a significant association are presented in Table 5.

**Figure 5:**
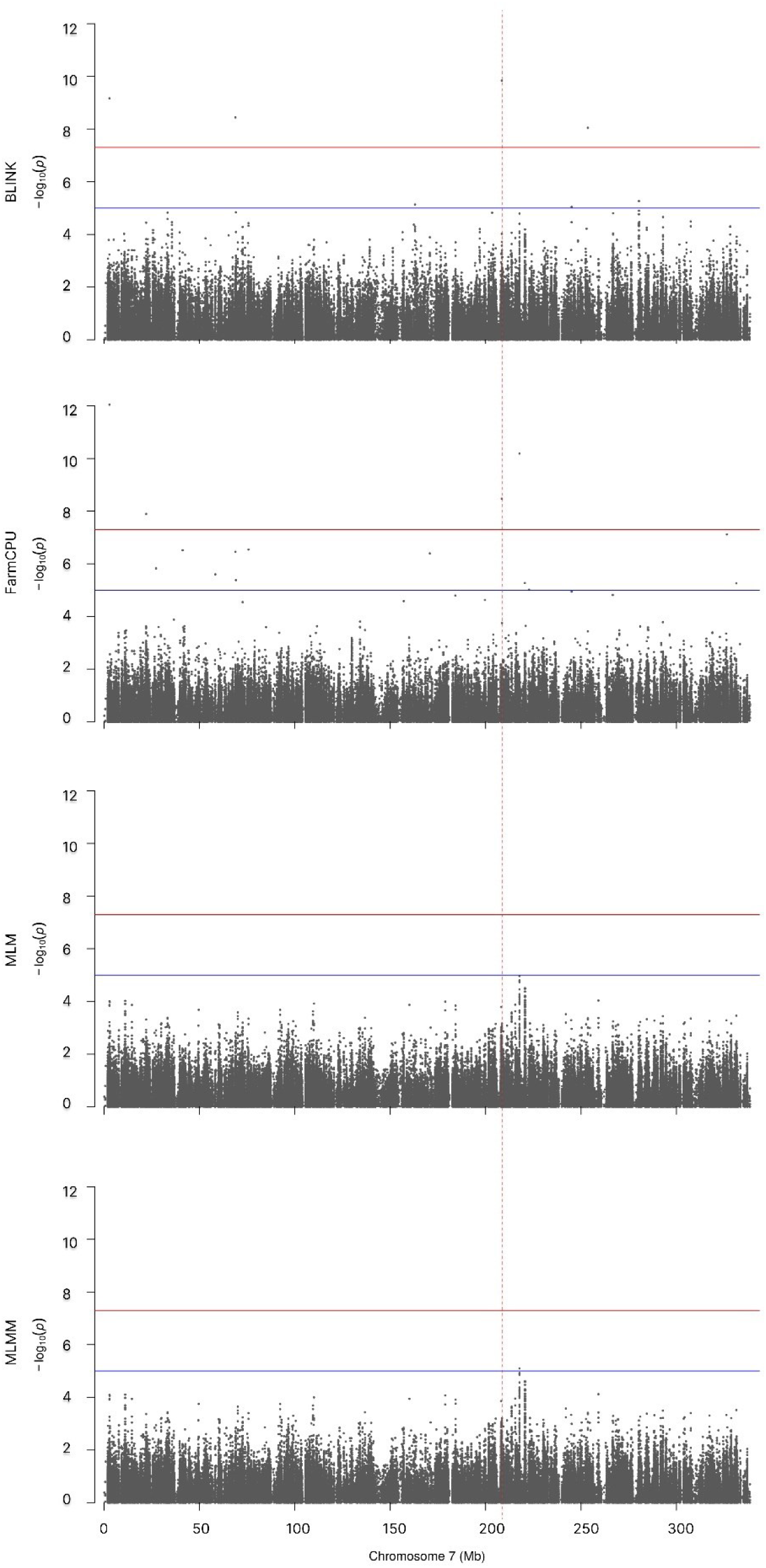
Genome-wide association analysis for seed shattering (%) on chromosome 7 using BLINK, FarmCPU, MLM and MLMM. Associations that were significant with the BLINK and/or FarmCPU model and showed a peak in the Manhattan plot using the MLM and MLMM models were highlighted with a red dashed vertical line. The red horizontal line indicates the Bonferroni-corrected threshold of 5%. The blue horizontal line indicates a *p*-value of *p* = 1.0 × 10^-5^.

**Table 4.** Single nucleotide polymorphisms (SNPs) associated with seed shattering (%), seed yield per plant, flowering time (days from first of April), spike length (cm), spike length (cm), spikelets per spike and flag leaf length (cm) in a nested association mapping population of Italian ryegrass (Lolium multiflorum Lam.). Chr indicates the chromosome, the position for each marker is shown in base pairs (bp). Gene name represents the name of the gene as annotated in the reference genome of cv. ’Rabiosà (Chen et al., 2025). The gene functions were revealed using the basic local alignment search tool (BLASTP) of NCBI.

| Trait | Chr | Marker position (bp) | Allelic effect | Phenotypic variance explained (%) | Gene name | Gene function |
| --- | --- | --- | --- | --- | --- | --- |
| Seed shattering (%) | 7 | 208,458,663 | 0.09 | 10.03 | *Chr7.26897* | Putative ripening-related protein 5 |
| Seed yield per plant (g) | 1 | 112,230,539 | -2.06 | 9.59 | - | Intergenetic region |
| Flowering time (d) | 4 | 57,744,424 | -1.83 | 33.59 | - | Intergenetic region |
|  | 4 | 95,647,826 | -1.07 | 10.72 | - | Intergenetic region |
|  | 6 | 113,373,604 | -1.46 | 16.75 | - | Intergenetic region |
| Spike length (cm) | 6 | 162,392,838 | -5.54 | 39.82 | - | Intergenetic region |
| Flag leaf length (cm) | 4 | 252,294,661 | 1.18 | 11.94 |  | Intergenetic region |

**Table 5.** Putative candidate genes identified for seed shattering (%), seed yield per plant (g), flowering time (start flowering counting from the 1sth of April), spike length (cm), number of spikelets per spike and flag leaf length (cm) using genome-wide association studies in the nested association mapping population of Italian ryegrass (*Lolium multiflorum* Lam.). Chr indicates the chromosome, the position for each gene is shown in base pairs. Gene name represents the name of the gene as annotated in the reference genome of cv. ’Rabiosà (Chen et al., 2025). The putative gene functions were revealed by using the basic local alignment search tool of NCBI.

| Trait | Chr | Position (bp) | Gene name | Putative gene function |
| --- | --- | --- | --- | --- |
| Seed shattering (%) | 7 | 208,458,401-208,458,868 | *Chr7.26897* | Putative ripening-related protein 5 |
| Seed yield per plant (g) | 1 | 112,223,339-112,237,739 | - | No gene annotated |
| Flowering time | 4 | 57,741,528-57,742,226 | *Chr4.7893* | PRA1 family protein |
|  | 4 | 95,652,729-95,655,604 | *Chr4.12997* | Unknown |
|  | 4 | 95,659,446-95,662,715 | *Chr4.12999* | Unknown |
|  | 6 | 113,366,404-113,380,804 | - | No gene annotated |
| Spike length (cm) | 6 | 162,385,638-162,400,038 | - | No gene annotated |
| Flag leaf length (cm) | 4 | 252,287,461-252,301,861 |  | No gene annotated |

For seed shattering, one significant association was found on chromosome 7 (Figure 5), which explained 10.03% of the phenotypic variance. This A/G SNP at position 208,458,663 bp was located inside the gene *Chr7.26897,* which has a function as a ripening-related protein.

The marker effect of this SNP was 0.05%. (Supplementary Figure S7a). Within the region of 14 kb up- and downstream of position 208,458,663 bp no other genes were annotated.

For seed yield per plant, one SNP significantly associated with the trait was found on chromosome 1 (Figure 6). This SNP on position 112,230,539 bp explained 9.59% of the phenotypic variance. The marker effect of this SNP was 2.01 g. Heterozygous plants (GA) at this position showed the highest seed yield per plant (Supplementary Figure S7b). No gene was found within the region of 14.40 kb up- and downstream of position 112,230,539 bp.

**Figure 6:**
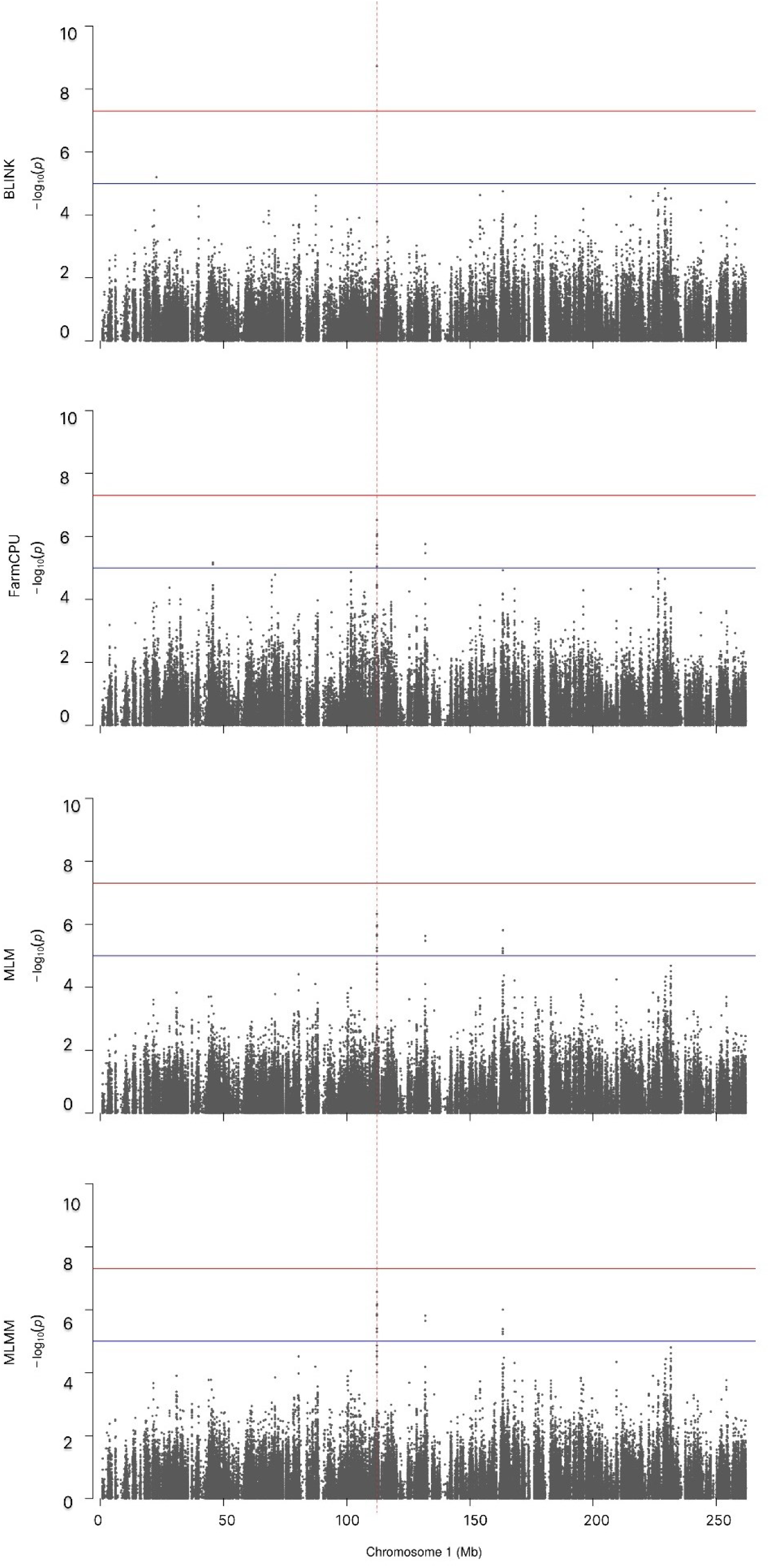
Genome-wide association analysis for seed yield per plant (g) on chromosome 1 using BLINK, FarmCPU, MLM and MLMM. Associations that were significant with the BLINK and/or FarmCPU model and showed a peak in the Manhattan plot using the MLM and MLMM models were highlighted with a red dashed vertical line. The red horizontal line indicates the Bonferroni-corrected threshold of 5%. The blue horizontal line indicates a *p*-value of *p* = 1.0 × 10^-5^.

For flowering time, one QTL was detected on chromosome 4 (Figure 7) and another on chromosome 6 (Figure 8). On chromosome 4, the two SNPs significantly associated with flowering time, located at positions 57,744,424 bp and 95,647,826 bp, explained 33.59% and 10.72% of the phenotypic variation, respectively. The marker effect of this SNP was 1.80 d. Plants with allele A at SNP position 57,744,424 bp were found to have on average a 1.80 d delay in flowering time compared to plants with allele G. Heterozygous plants (GA) showed the largest delay in flowering time compared to homozygous AA and GG plants (Supplementary, Figure S7c). Plants with allele A at SNP position 95,647,826 bp starting on average 0.83 d later to flower. Heterozygous plants (GA) at this marker position flowered the latest (Supplementary, Figure S7d). Both SNP positions are not within an annotated gene. However, within 14.40 kb up- and downstream from marker 57,744,424 bp, one gene was annotated a PRA1 family protein (*Chr4.7893*). For the significant marker-trait association on position 95,647,826, two genes were annotated within a region of 14.4 kb up- and downstream, both with unknown functions. On chromosome 6, the marker most significantly associated with flowering time on position 113,373,604 bp explained 16.75% of the phenotypic variance. Plants with allele G at this position showed on average a 1.33 d later flowering compared to the plants with allele A. Plants being heterozygous at this position showed the latest flowering time compared to the rest of the plants (Supplementary Figure S7e).

**Figure 7:**
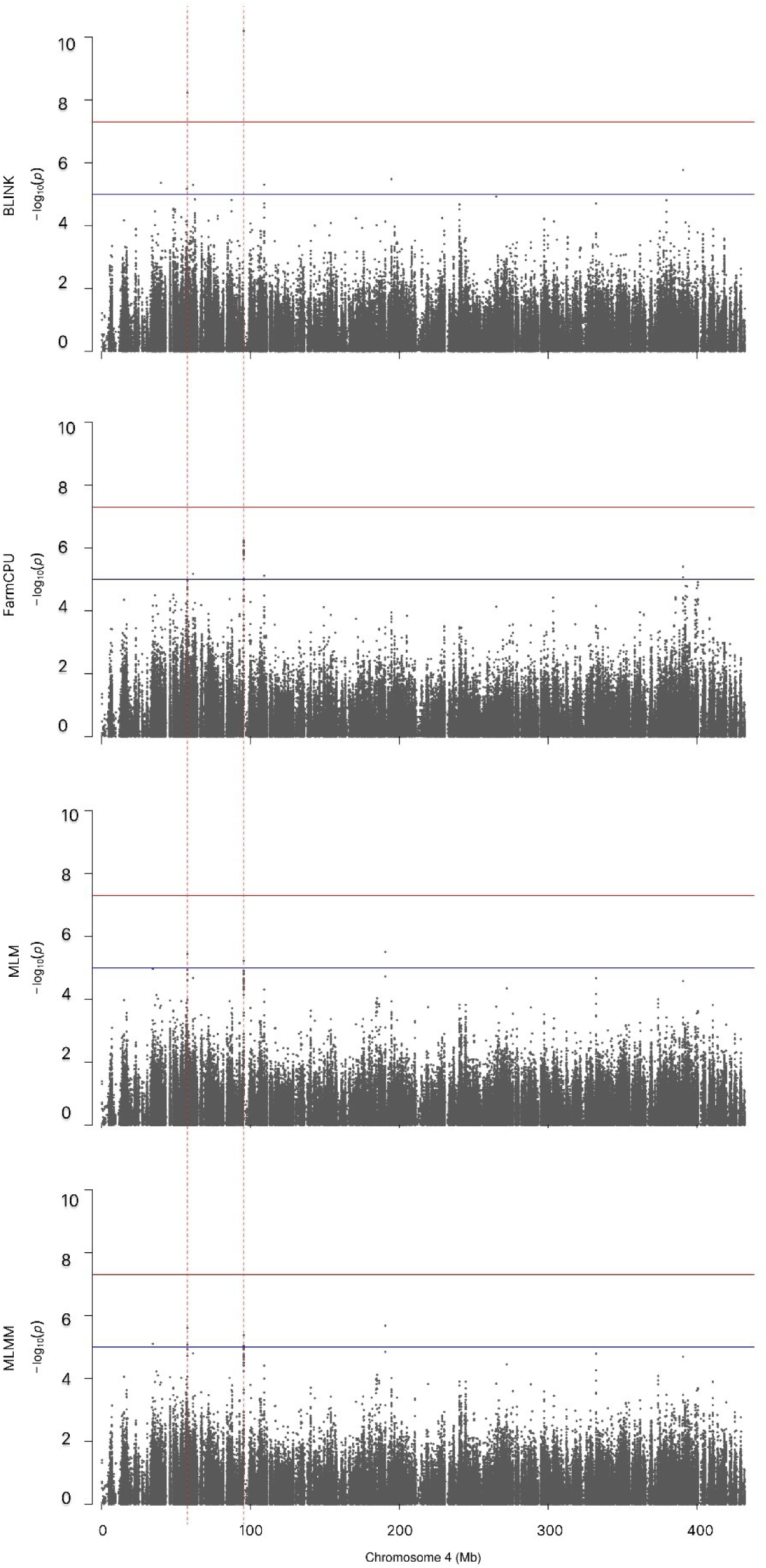
Genome-wide association analysis for flowering time on chromosome 4 using BLINK, FarmCPU, MLM and MLMM. Associations that were significant with the BLINK and/or FarmCPU model and showed a peak in the Manhattan plot using the MLM and MLMM models were highlighted with a red dashed vertical line. The red horizontal line indicates the Bonferroni-corrected threshold of 5%. The blue horizontal line indicates a *p*-value of *p* = 1.0 × 10^-5^.

**Figure 8:**
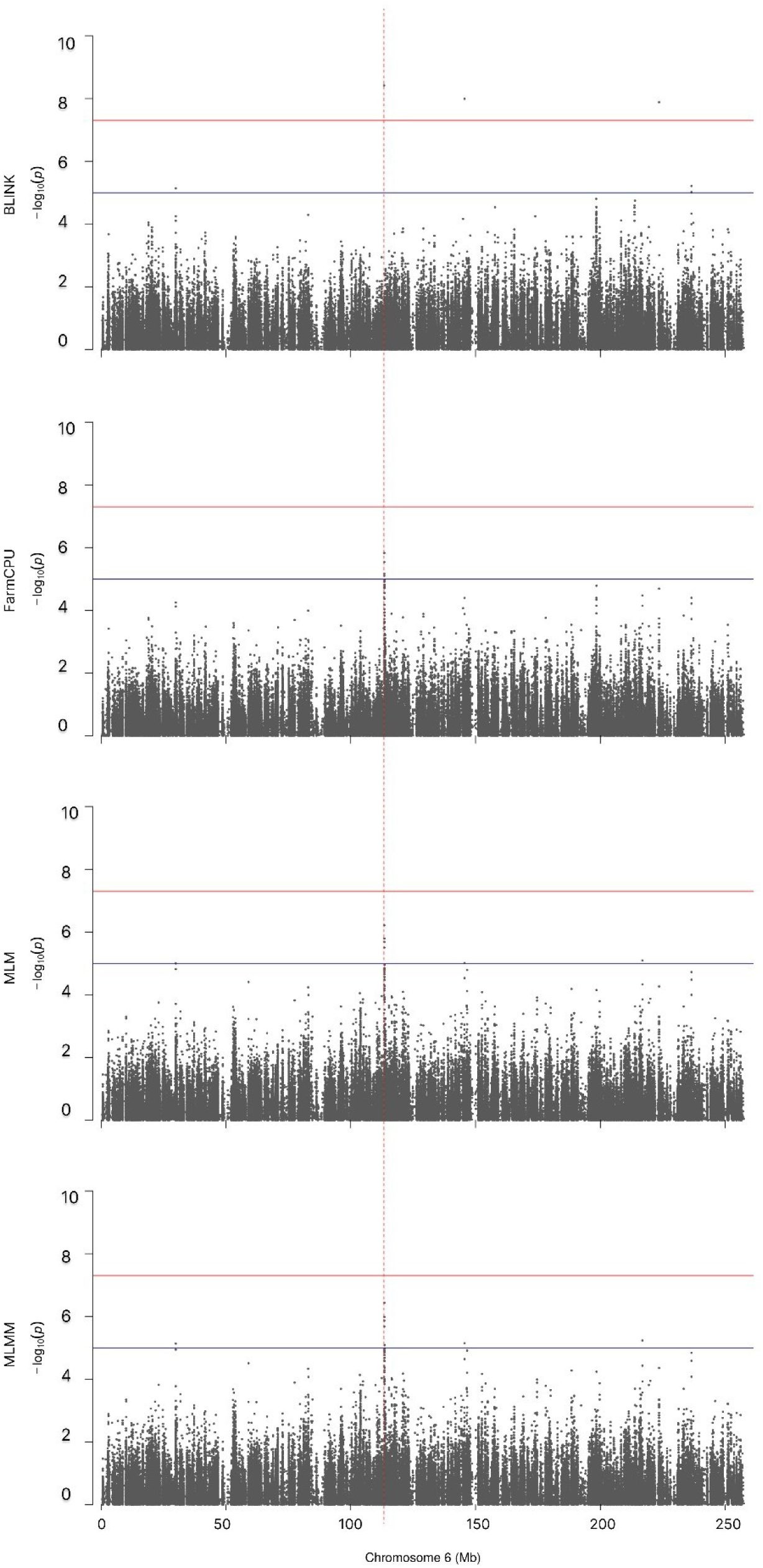
Genome-wide association analysis for flowering time on chromosome 6 using BLINK, FarmCPU, MLM and MLMM. Associations that were significant with the BLINK and/or FarmCPU model and showed a peak in the Manhattan plot using the MLM and MLMM models were highlighted with a red dashed vertical line. The red horizontal line indicates the Bonferroni-corrected threshold of 5%. The blue horizontal line indicates a *p*-value of *p* = 1.0 × 10^-5^.

One SNP on chromosome 6 on position 162,392,838 bp showed a significant association with spike length (Figure 9). This SNP explained 39.82% of the phenotypic variation. Plants with allele G showed on average 5.11 cm larger spikes compared to plants with allele A at this position. Heterozygous plants (GA) at this position showed the largest spikes (Supplementary Figure S7f). No gene was annotated within 14.40 kb up-downstream of the significantly associated SNP. No significant associations were found for the trait spikes per spikelet (Figure 10).

**Figure 9:**
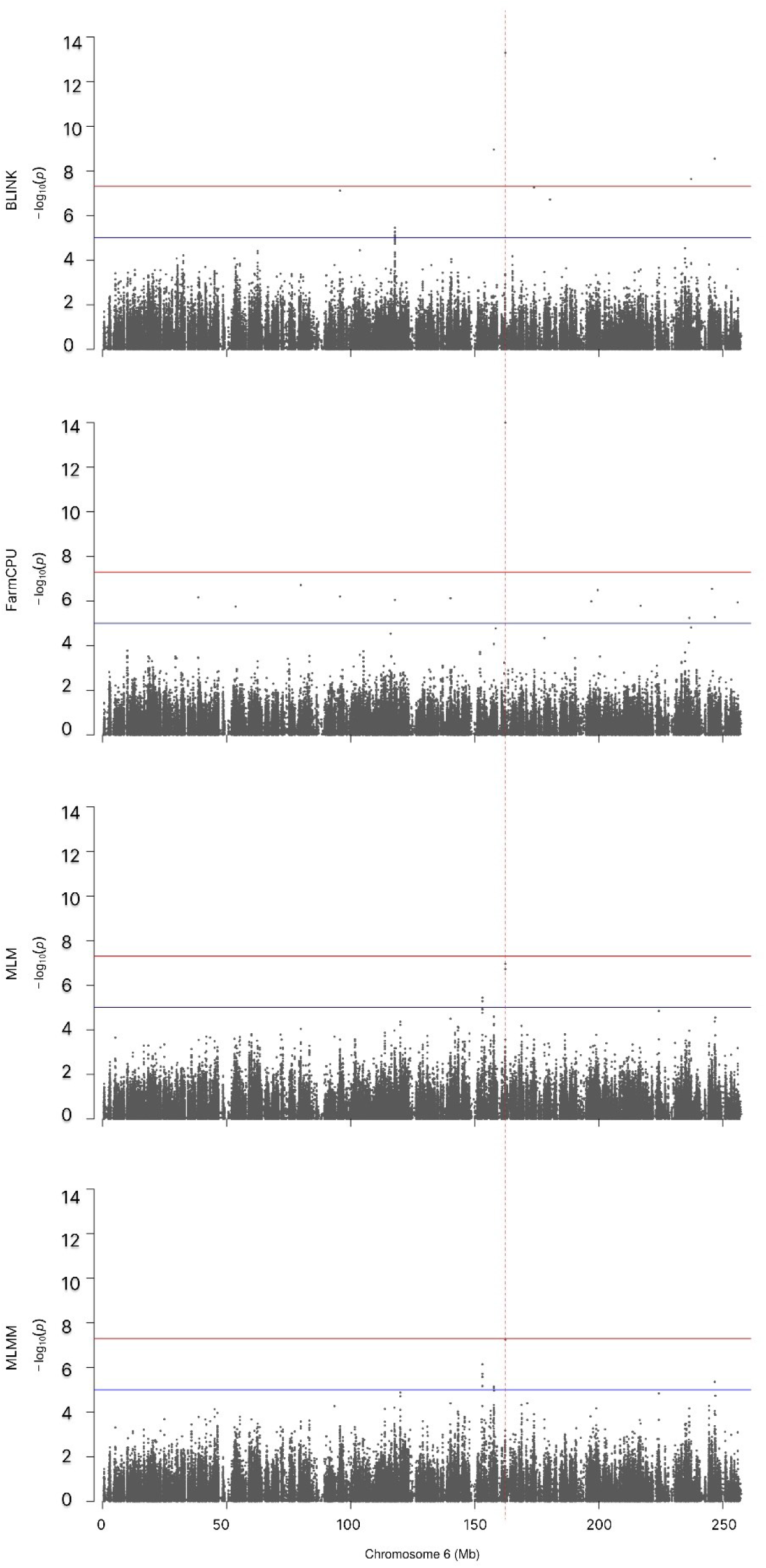
Genome-wide association analysis for spike length (cm) on chromosome 6 using BLINK, FarmCPU, MLM and MLMM. Associations that were significant with the BLINK and/or FarmCPU model and showed a peak in the Manhattan plot using the MLM and MLMM models were highlighted with a red dashed vertical line. The red horizontal line indicates the Bonferroni-corrected threshold of 5%. The blue horizontal line indicates a *p*-value of *p* = 1.0 × 10^-5^.

**Figure 10:**
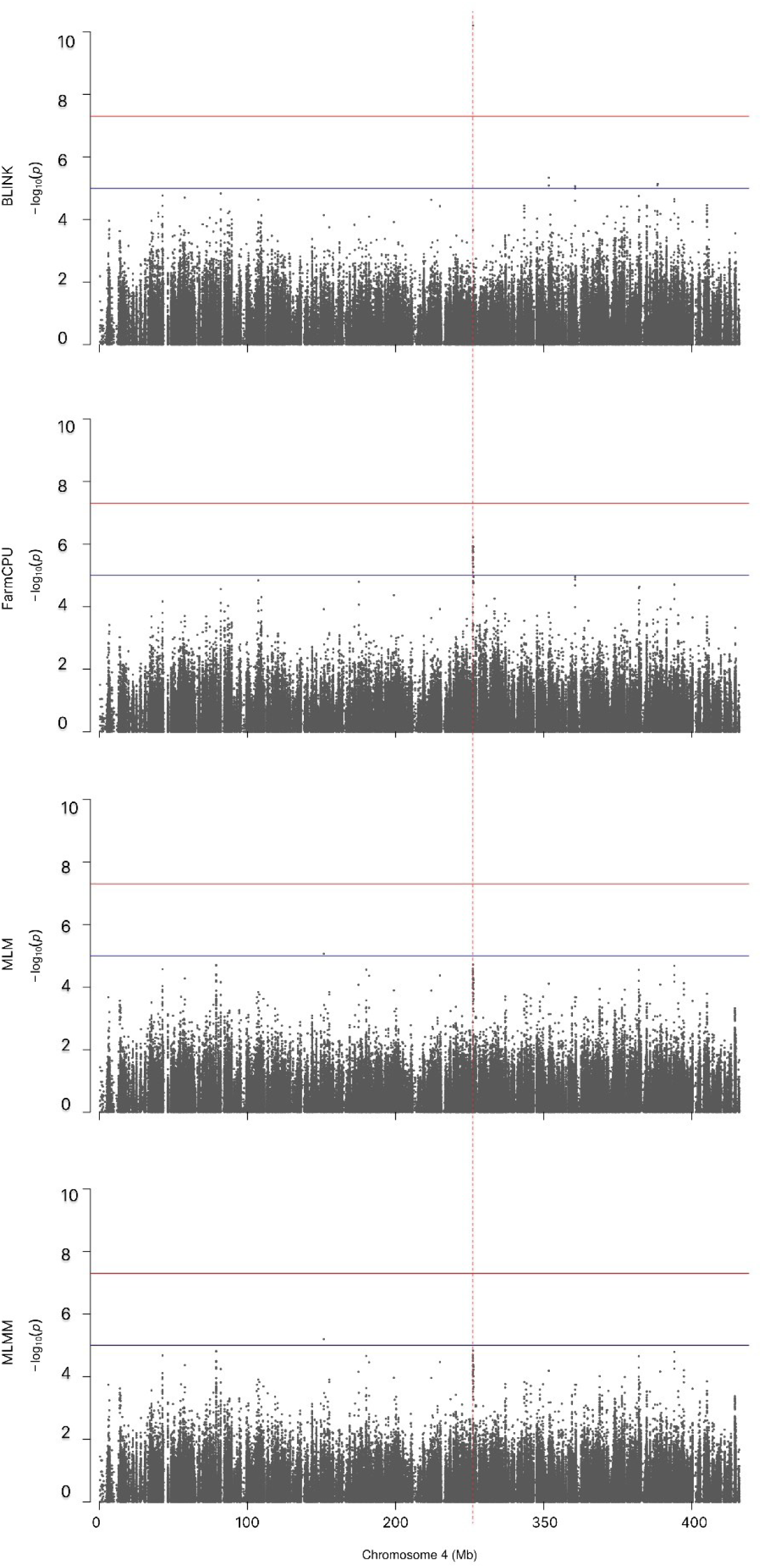
Genome-wide association analysis for flag leaf length (cm) on chromosome 4 using BLINK, FarmCPU, MLM and MLMM. Associations that were significant with the BLINK and/or FarmCPU model and showed a peak in the Manhattan plot using the MLM and MLMM models were highlighted with a red dashed vertical line. The red horizontal line indicates the Bonferroni-corrected threshold of 5%. The blue horizontal line indicates a *p*-value of *p* = 1.0 × 10^-5^.

One SNP significantly associated with flag leaf length on chromosome 4 at position 252,294,661 bp explained 11.94% of the phenotypic variance. Plants with allele G at this position showed on average 0.94 cm longer flag leaves compared to plants with allele A. Heterozygous plants showed the lowest flag leaf length (Supplementary Figure S7g). Within 14.40 kb up- and downstream from the significantly associated SNP, no gene was annotated.

## Discussion

This study presents the first NAM population developed for Italian ryegrass, an agronomically important outcrossing forage grass species. The NAM population consisted of 708 F_2_ individuals and was designed to account for crossing-over events over two generations, both of which contribute to increased mapping resolution. The design included a common founder, with a chromosome-scale, haplotype-resolved genome sequence recently published (Chen et al., 2025). The design also incorporated 23 diverse founder plants, selected for high phenotypic diversity in general and for high variation in seed shattering in particular. Further, to ensure statistical detection power, a crossing scheme with a balanced family structure and therefore a minimal allele frequency bias was used. Another key feature of the NAM is the integration of high-coverage WGS data of all founders combined with ddRAD data from the F₂ individuals, enabling SNP imputation at highest possible resolution. The utility of the NAM population was validated through GWAS and candidate gene identification on several genetically complex and economically important traits, including seed shattering, flowering time and seed yield. This demonstrates the robustness and precision of this resource for trait dissection and sets a solid foundation for genomics-assisted breeding in Italian ryegrass.

To date, NAM populations in major crops were mainly designed based on pairwise crossings of one common parent with many diverse donor parents, followed by repeated self-fertilization to produce RILs (Yu et al., 2008). However, this mating scheme is not applicable for outcrossing species such as Italian ryegrass with high self-incompatibility and severely suffering from inbreeding depression. One NAM population for an obligate outcrossing species (intermediate wheat grass [*Thinopyrum intermedium*]) has previously been described for seed shattering and threshability (Altendorf et al., 2021). In that study, the NAM population consisted of a cross between one common founder and ten diverse founders, resulting in ten full-sib families (F_1_) related to the common founder (Altendorf et al., 2021). In contrast to our study, where the 23 F_1_ families were advanced to F_2_ by open pollination, no further crosses were made and informative recombination events from 1,168 F_1_ progeny were exploited in that study. The self-incompatibility system of Italian ryegrass is gametophytically controlled by two unlinked multi-allelic loci, known as *S* and *Z* (Cornish et al., 1979; Hayman, 1956; Lundqvist, 1954). Therefore, the system also prevents fertilization between two plants containing the same *S* and *Z* alleles, potentially having influenced the open pollination among the F_1_, resulting in a violation of the assumption of panmixia. However, no population structure was observed with respect to the founder plants, leading to the conclusion that there was no unequal pollination (Figure 2). This was expected, as the high number of founder plants resulted in high allele diversity at the S and Z loci. However, to ensure uniform pollination of the population, the founder plants were selected on the basis of high *S* and *Z* allele diversity (Rohner, 2022).

The population structure of the F_2_ NAM population was further investigated to reveal to what extent each founder contributed to the population. Even though the correlation analysis was able to identify more parents correctly, the two complementary approaches (KING software and correlation analysis) together form a robust basis for identifying the parents of each plant in the NAM population. Both approaches showed that all founders contributed to the open pollinated F_2_ NAM population. Most founders contributed equally to the NAM F_2_ population. However, F_1_ plants derived from either a cross between MD1, MD4, MD5, or MD20 with MC contributed less to population pollination (Table 1). One possible explanation is that the four F_1_ plants had a slightly different flowering time than the rest of the population. For the F_2_ crossings, tillers were used which can be usually put in the cold to delay flowering. Although attempts were made to synchronize the different flowering times through cold storage, this was apparently not always fully successful. However, the unequal distribution of a few founder plants seemed to have only a weak influence on the NAM F_2_ population structure.

High-quality phenotypic data are a prerequisite for identifying the genetic control underlying the respective trait. Our phenotypic data revealed moderate to high or even very high heritability (*h^2^* > 0.68) for all traits measured. High heritability results in a high statistical power to detect QTL with even small effects (Yu et al., 2008). This makes the generated phenotypic dataset a valuable resource also for the discovery of QTL such as seed yield, which is described as a highly quantitative and complex trait (Jaškūnė et al., 2022).

The features of our NAM population, combined with high-quality phenotypic data, now enabled a more precise dissection of the traits that determine seed yield. For the number of spikelets per spike, an important seed yield component, mainly influencing the potential seed yield, the mean value of our NAM F_2_ population was 18.49. This is lower in comparison to other studies in perennial ryegrass, where the mean value of the number of spikelets per spike was 22.00 (Chastain et al., 2014), 24.60 (Elgersma & Śnieżko, 1988), 19.30-28.10 (Tubbs & Chastain, 2022) and 24.20 (Yamada et al., 2004). This can be explained by the fact that the NAM F_2_ population is based on several ecotypes, which are often reported to have a lower number of spikelets per spike (McGrath et al., 2010). The phenotypic variation of spikelets per spike in the F_2_ NAM population was comparable or somewhat lower than in studies with perennial ryegrass (McGrath et al., 2010; Tubbs & Chastain, 2022). On the other hand, the mean value for spike length was higher in the F_2_ NAM population compared to previous studies in perennial ryegrass (Byrne et al., 2009; Tubbs & Chastain, 2022). Previous studies showed a correlation of 0.78 (Abel et al., 2017) and 0.46 (Byrne et al., 2009) between the number of spikelets per spike and spike length. These results are similar to the correlation of *r* = 0.50 observed here. Abel et al. (2017) suggest maximizing seed yield by increasing the spike length and, therefore, increasing the number of spikelets per spike. However, our study showed only weak to moderate correlation between seed yield and spike length (*r_p_* = 0.32) and breeding for longer spikes may therefore not be a promising strategy to achieve higher seed yield in Italian ryegrass. Elgersma et al. (1988) showed a strong correlation between harvestable seed yield and seed shattering. The hypothesis was, that plants with high seed numbers (e.g., high yielding plants) are prone to shattering due to the increased weight. In contrast to that study, only a very weak correlation between total seed yield (g) and seed shattering (%) was found within our F_2_ NAM population (*r_p_* = 0.11). Furthermore, high seed yielding cultivars seem not *per se* be more prone to seed shattering (Kiesbauer et al., 2025). Therefore, breeding a low seed shattering cultivar should be achievable without compromising other important agronomic traits such as total seed yield or flowering time.

The genetic basis for seed yield and seed yield-related traits is largely unknown in forage grasses. Previous QTL studies in perennial ryegrass revealed several loci associated with spikelets per spike, spike length, flag leaf length and seed yield (Byrne et al., 2009; Jaškūnė et al., 2022). A recent study showed a major QTL on linkage group 2 for reduced seed shattering, which appears to be partially dominantly inherited in Italian ryegrass (Tamura et al., 2024). In our study, a QTL on chromosome 7 was found that explained 10% of the phenotypic variance of seed shattering. The significant SNP was located within a gene (*Chr7.26897*) with a ripening-related function (BLASTP 100% identity to *L. rigidum* ripening-related protein (*LOC124655758*)). Early and late timing of harvesting were tested for their influence on seed shattering in the same genotype of single plants in Italian ryegrass, whereby early harvesting reduced the percentage of seed shattering (Kiesbauer et al., 2025). Therefore, ripening and thus the putative ripening related gene which was found here may play a role in seed shattering.

For flowering time, several studies identified more than 30 QTL across all chromosomes in ryegrass (Armstead et al., 2004, 2008; Barre et al., 2009; Byrne et al., 2009; Jensen et al., 2005; King et al., 2008; Studer et al., 2008; Wang et al., 2011; Yamada et al., 2004), some consistently detected across the different studies (Armstead et al., 2009, 2008). One known flowering gene, *LpFT3,* is located on chromosome 7 in perennial ryegrass (Skøt et al., 2010). However, our candidate genes, located on chromosome 4 and 6, have not been previously described to have a known function in flowering time in ryegrass (Wang & Forster, 2017). Not only for flowering time, but also for all the other traits measured, none of our results are consistent with previously published QTL. A possible explanation is that the genomic resources such as high-quality reference genomes and sequencing technologies have only recently become available and affordable for ryegrass. Previously, the main approach to locating loci associated with traits of interest in ryegrass was based mainly on biparental mapping populations, resulting in very specific results for the investigated population, making them difficult to transfer to other populations. In comparison to a biparental population, a NAM population design can capture diverse as well as enrich rare alleles for traits of interest and therefore minor QTL are detectable. Another possible explanation could be the highly quantitative nature of these traits, i.e., they are each slightly influenced by many different genes and therefore specific to particular populations. Further, many biparental populations were established based on parents which are only interesting for one particular trait but lacking a suitable agronomic performance and therefore can not be easily included in the conventional breeding program. Alternatively, with a multi-parental mapping population, such as a NAM population, different breeding material can be easily included, and the results are transferable to breeding. For example, in our NAM population, 14 out of 23 diverse founders were selected from breeding material and the results can be incorporated directly into the breeding program using genomic-assisted selection.

To conclude, we successfully established a NAM F_2_ population in Italian ryegrass. Our NAM population design led to almost no population structure which would potentially negatively influence QTL mapping. The sequencing strategy of sequencing the 23 diverse founders by whole genome sequencing, followed by ddRAD sequencing the NAM F_2_ population is a very cost-effective strategy which allowed for a high density of imputed SNPs in the NAM F_2_ population. Together with high-quality phenotypic data, eight novel loci could be identified for seed shattering, seed yield, flowering time, spike length and flag leaf length. As many NAM founders are breeding material, the SNPs associated with the trait of interest can be directly used for genomic-assisted selection. For seed shattering as well as for flowering time, we were able to identify four candidate genes. However, it is also possible that the association between trait and marker reflect regulatory or non-coding variation. Therefore, the listed candidate genes need to be functionally validated in upcoming research projects. The outcome of this study highlights the value of establishing a NAM population for the dissection of complex traits such as seed yield and related traits in Italian ryegrass.

## Data availability

The mapped WGS and ddRAD files are available in the NCBI Sequence Read Archive as BioProject accession number PRJNA1167222.

## Acknowledgements

Genotyping data of the NAM F_2_ population were generated in collaboration with the Genetic Diversity Centre (GDC), ETH Zurich. We thank the Functional Genomic Centre Zurich (FGCZ) for sequencing the NAM founders. This project was funded by the Breeding Foundation DSP-BLW

## Supplementary

**Table S1:** Name and description of the accessions from which the common (M0) and diverse founder plants (M1-23) used for the establishment of the F2 nested association mapping population for Italian ryegrass *(Lolium multiflorum* Lam.) were selected from.

| M | Name | Description |
| --- | --- | --- |
| 0 | Rabiosa | Cultivar from fodder plant breeding group, division of plant breeding, Agroscope, Zurich, Switzerland |
| 1 | Sikem | Cultivar from DLF, Roskilde, Denmark |
| 2 | Shakira | Cultivar from DSV, Lippstadt, Germany |
| 3 | Srl1_LI1115_3501-02 | Breeding material from fodder plant breeding group, division of plant breeding, Agroscope, Zurich, Switzerland |
| 4 | Srl2_LI1266_3413-10 | Breeding material from fodder plant breeding group, division of plant breeding, Agroscope, Zurich, Switzerland |
| 5 | Srl3_LI1245_3426-10 | Breeding material from fodder plant breeding group, division of plant breeding, Agroscope, Zurich, Switzerland |
| 6 | LI9901 | Breeding material from fodder plant breeding group, division of plant breeding, Agroscope, Zurich, Switzerland |
| 7 | LI9902 | Breeding material from fodder plant breeding group, division of plant breeding, Agroscope, Zurich, Switzerland |
| 8 | Arth_Chraebel | https://www.pgrel.admin.ch/pgrel/#/objects/accessions/detail/0000042971/description |
| 9 | Cadenazzo | https://www.pgrel.admin.ch/pgrel/#/objects/accessions/detail/0000042952/description |
| 10 | Castel_S.Pietro | https://www.pgrel.admin.ch/pgrel/#/objects/accessions/detail/0000042949/description |
| 11 | Egg | https://www.pgrel.admin.ch/pgrel/#/objects/accessions/detail/0000039220/description |
| 12 | Tremona | https://www.pgrel.admin.ch/pgrel/#/objects/accessions/detail/42950/description |
| 13 | Davinci | Cultivar from DLF, Roskilde, Denmark |
| 14 | Majesty | Cultivar from DSV, Lippstadt, Germany |
| 15 | Oryx | Cultivar from fodder plant breeding group, division of plant breeding, Agroscope, Zurich, Switzerland |
| 16 | Rangifer | https://www.pgrel.admin.ch/pgrel/#/objects/accessions/detail/0000046816/description |
| 17 | LI0085 | Breeding material from fodder plant breeding group, division of plant breeding, Agroscope, Zurich, Switzerland |
| 18 | LI0505 | Breeding material from fodder plant breeding group, division of plant breeding, Agroscope, Zurich, Switzerland |
| 19 | LI0615 | Breeding material from fodder plant breeding group, division of plant breeding, Agroscope, Zurich, Switzerland |
| 20 | LI0905 | Breeding material from fodder plant breeding group, division of plant breeding, Agroscope, Zurich, Switzerland |
| 21 | LI0915 | Breeding material from fodder plant breeding group, division of plant breeding, Agroscope, Zurich, Switzerland |
| 22 | LI1015 | Breeding material from fodder plant breeding group, division of plant breeding, Agroscope, Zurich, Switzerland |
| 23 | LI1135 | Breeding material from fodder plant breeding group, division of plant breeding, Agroscope, Zurich, Switzerland |

**Figure S1.**
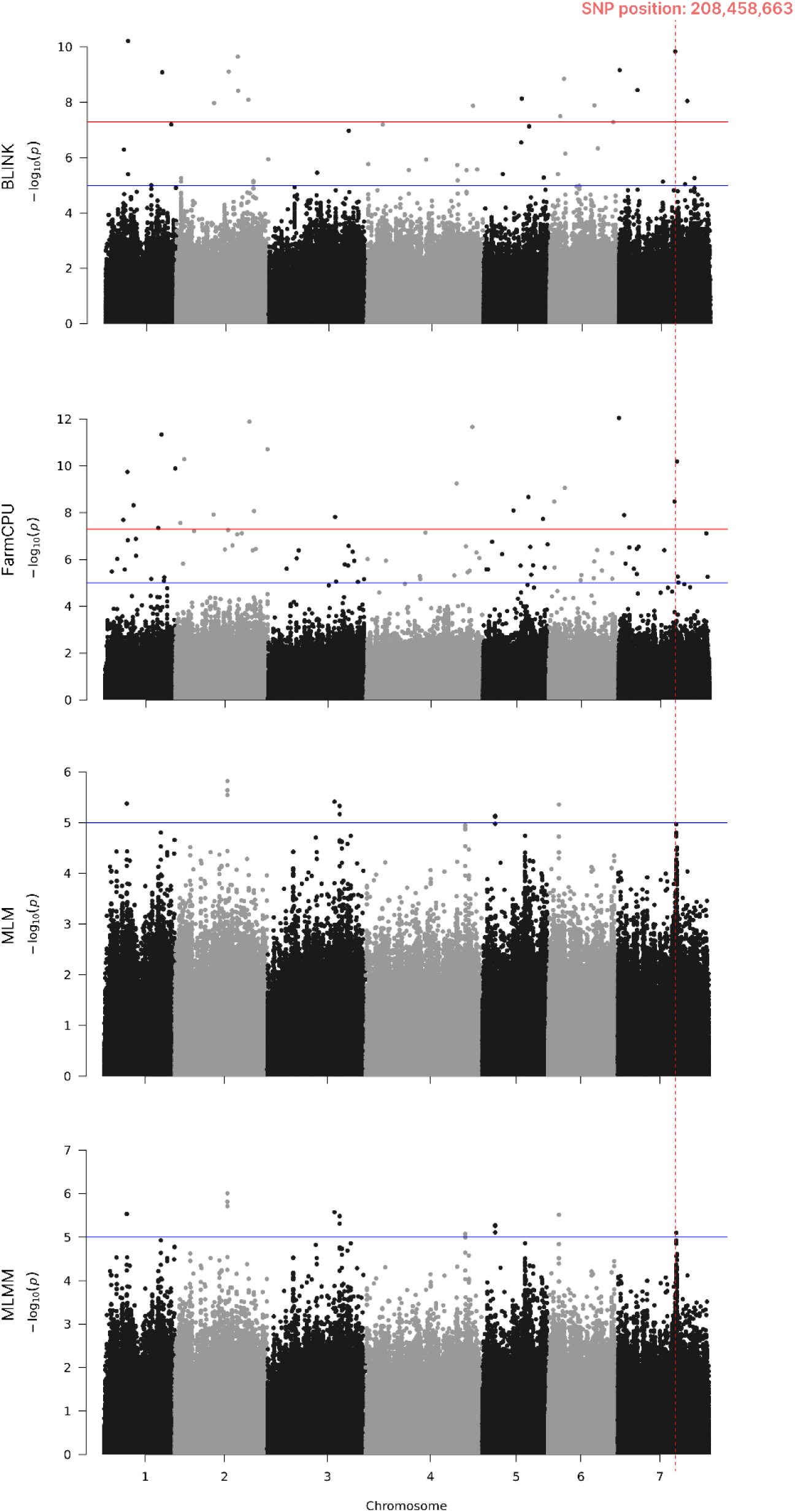
Genome-wide association analysis for **seed shattering** using BLINK, FarmCPU, MLM and MLMM. The dashed red vertical line indicates the common markers, which were found by all four methods being at least significantly associated by using BLINK. The red horizontal line indicates the Bonferroni-corrected threshold of 5%. The blue horizontal line indicates a *p-*value of *p* = 1.0 × 10^-5^.

**Figure S2.**
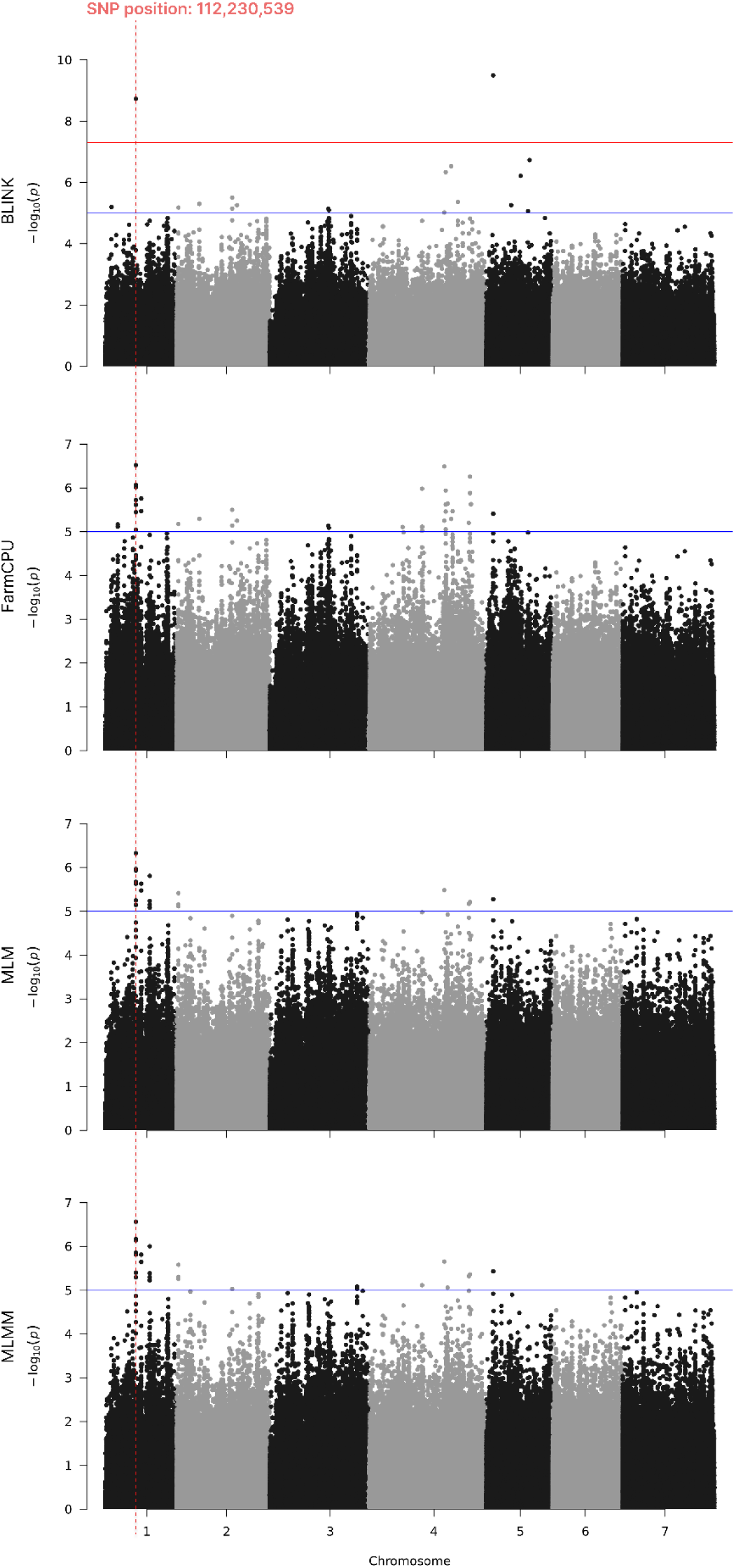
Genome-wide association analysis for **seed yield** using BLINK, FarmCPU, MLM and MLMM.. The dashed red vertical line indicates the common markers, which were found by all four methods being at least significantly associated by using BLINK. The red horizontal line indicates the Bonferroni-corrected threshold of 5%. The blue horizontal line indicates a *p*-value of *p* = 1.0 × 10^-5^.

**Figure S3.**
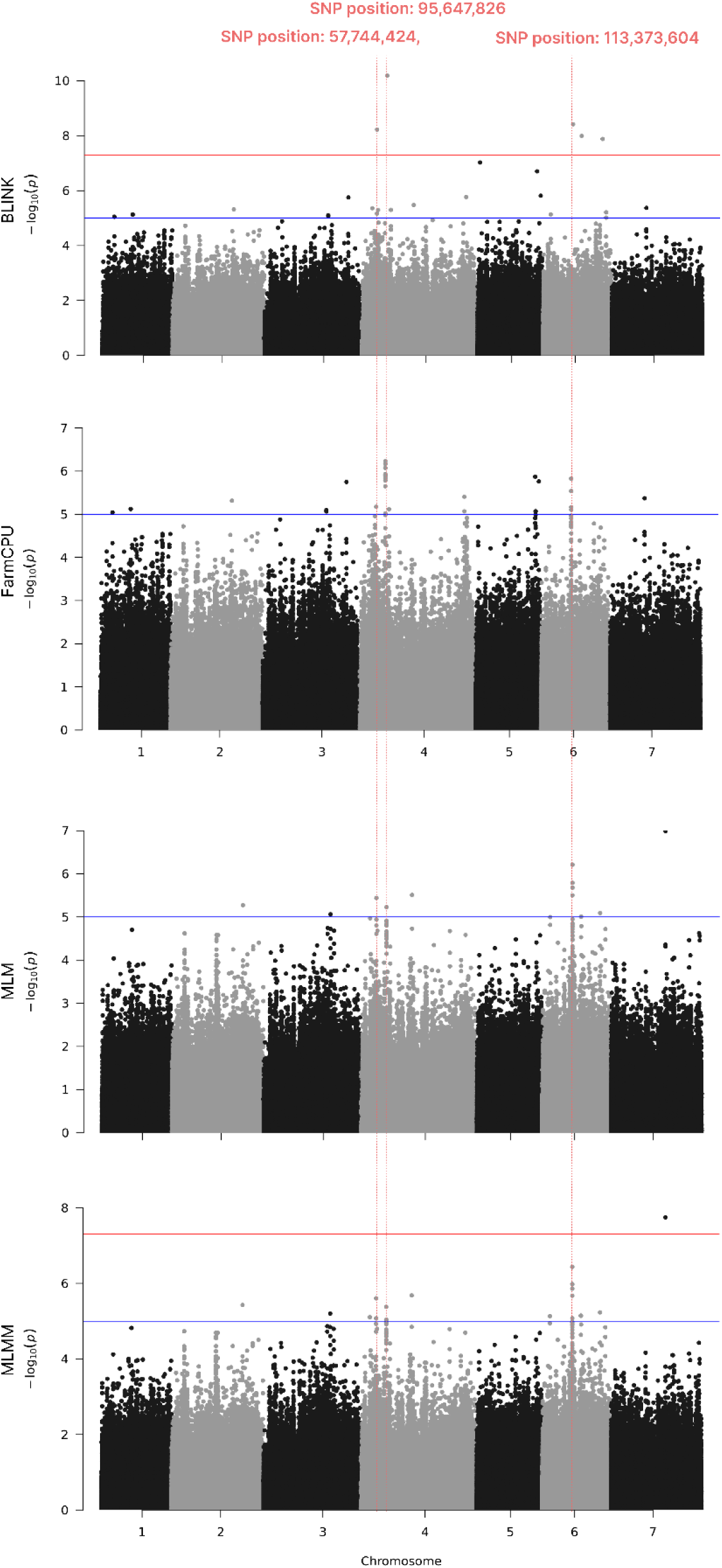
Genome-wide association analysis for **flowering time** using BLINK, FarmCPU, MLM and MLMM.. The dashed red vertical line indicates the common markers, which were found by all four methods being at least significantly associated by using BLINK. The red horizontal line indicates the Bonferroni-corrected threshold of 5%. The blue horizontal line indicates a *p*-value of *p* = 1.0 × 10^-5^.

**Figure S4.**
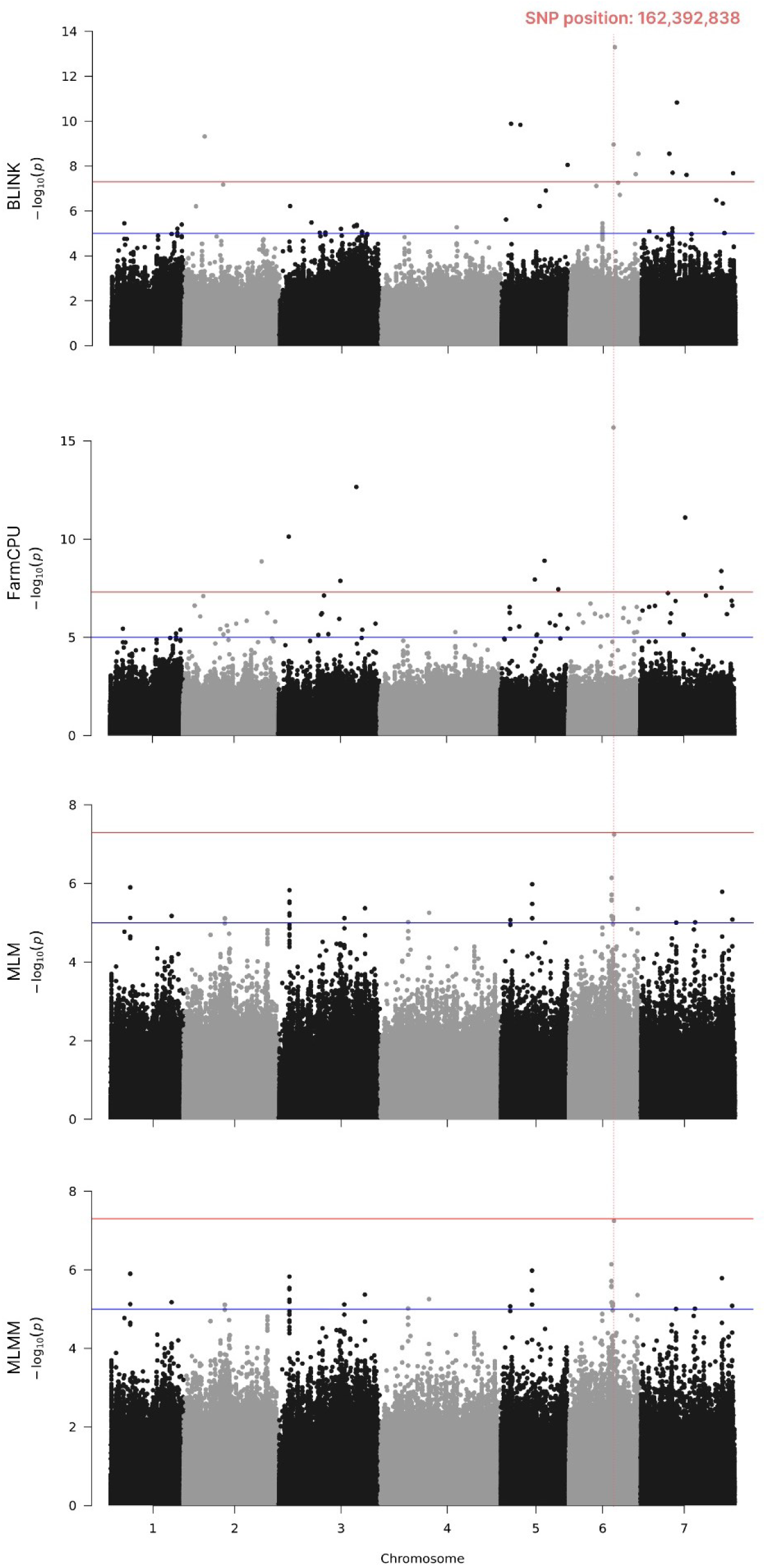
Genome-wide association analysis for **spike length** using BLINK, FarmCPU, MLM and MLMM. The dashed red vertical line indicates the common markers, which were found by all four methods being at least significantly associated by using BLINK. The red horizontal line indicates the Bonferroni-corrected threshold of 5%. The blue horizontal line indicates a *p*-value of *p* = 1.0 × 10^-5^.

**Figure S5.**
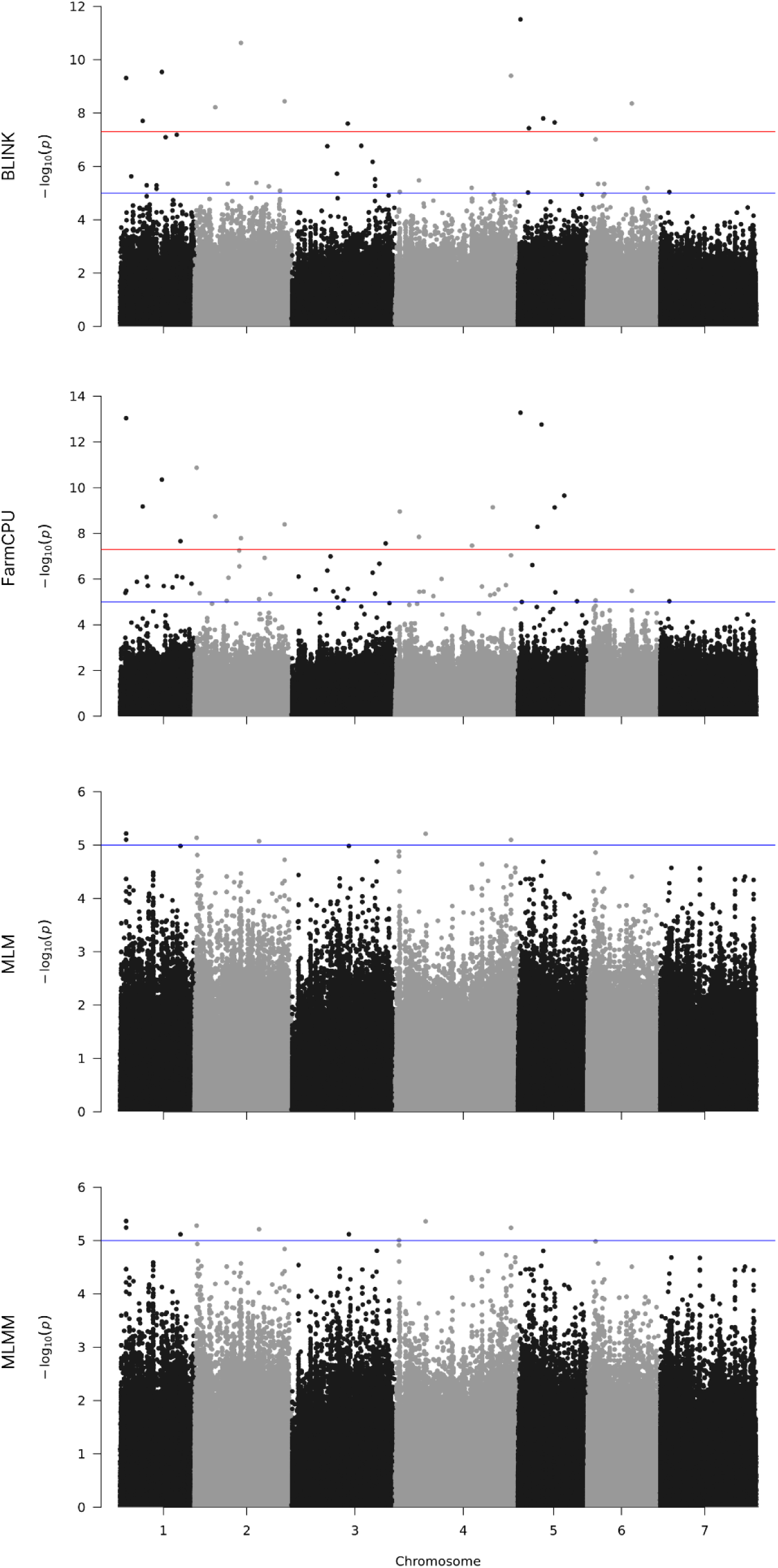
Genome-wide association analysis for **spikelets per spike** using BLINK, FarmCPU, MLM and MLMM. The dashed red vertical line indicates the common markers, which were found by all four methods being at least significantly associated by using BLINK. The red horizontal line indicates the Bonferroni-corrected threshold of 5%. The blue horizontal line a *p*-value of *p* = 1.0 × 10^-5^.

**Figure S6.**
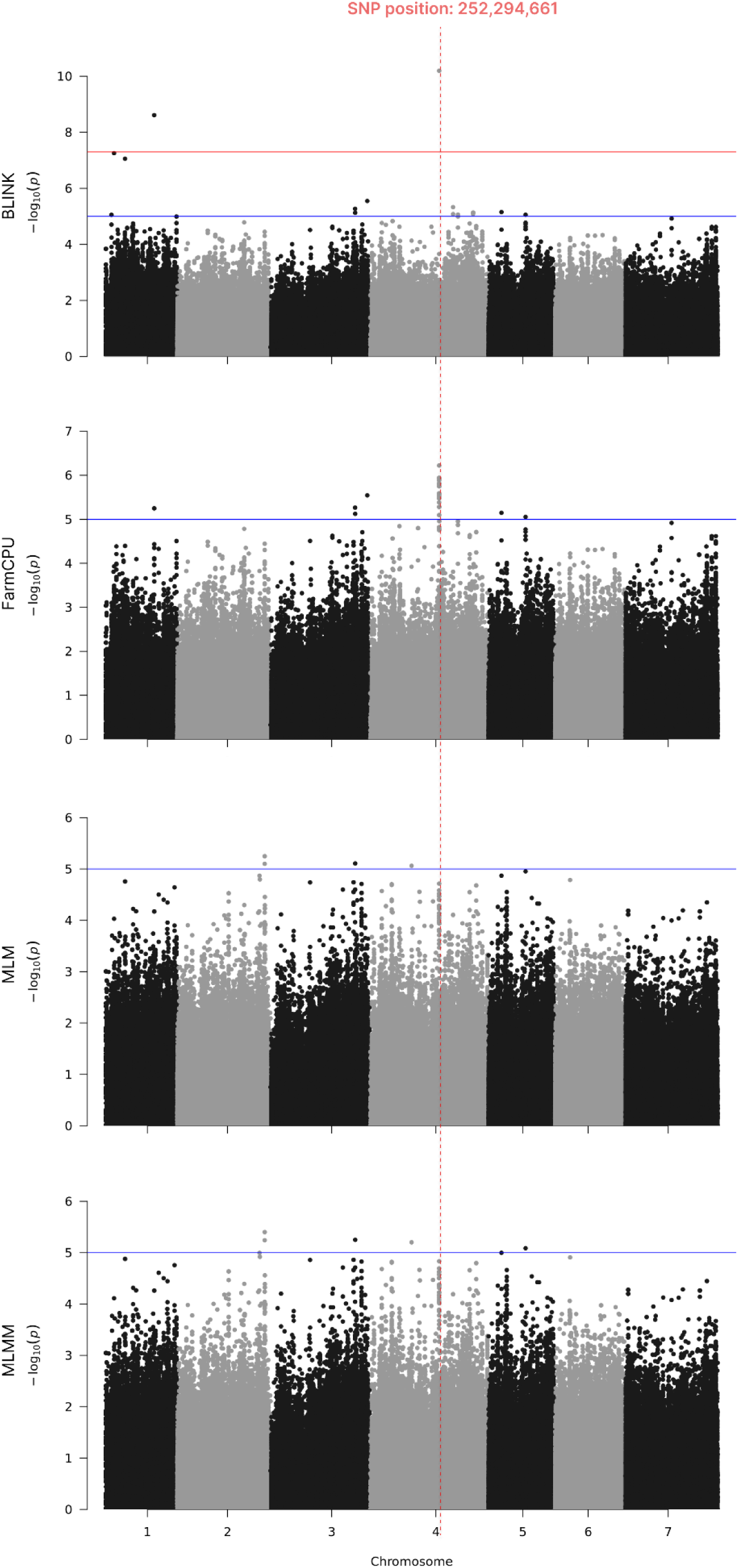
Genome-wide association analysis for **flag leaf length** using FarmCPU, BLINK, MLM and MLMM. The dashed red vertical line indicates the common markers, which were found by all four methods being at least significantly associated by using BLINK. The red horizontal line indicates the Bonferroni-corrected threshold of 5%. The blue horizontal line indicates a *p*-value of *p* = 1.0 × 10^-5^.

**Figure S7.**
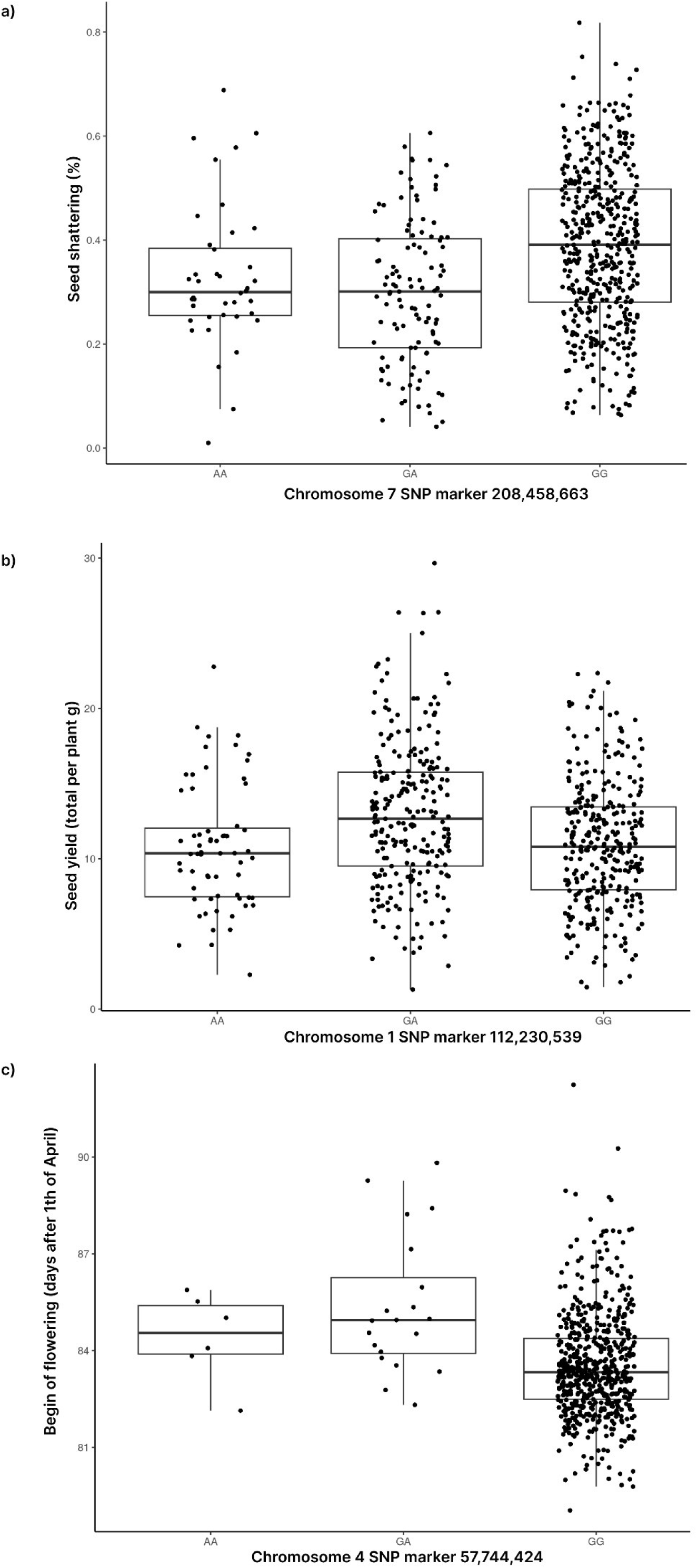
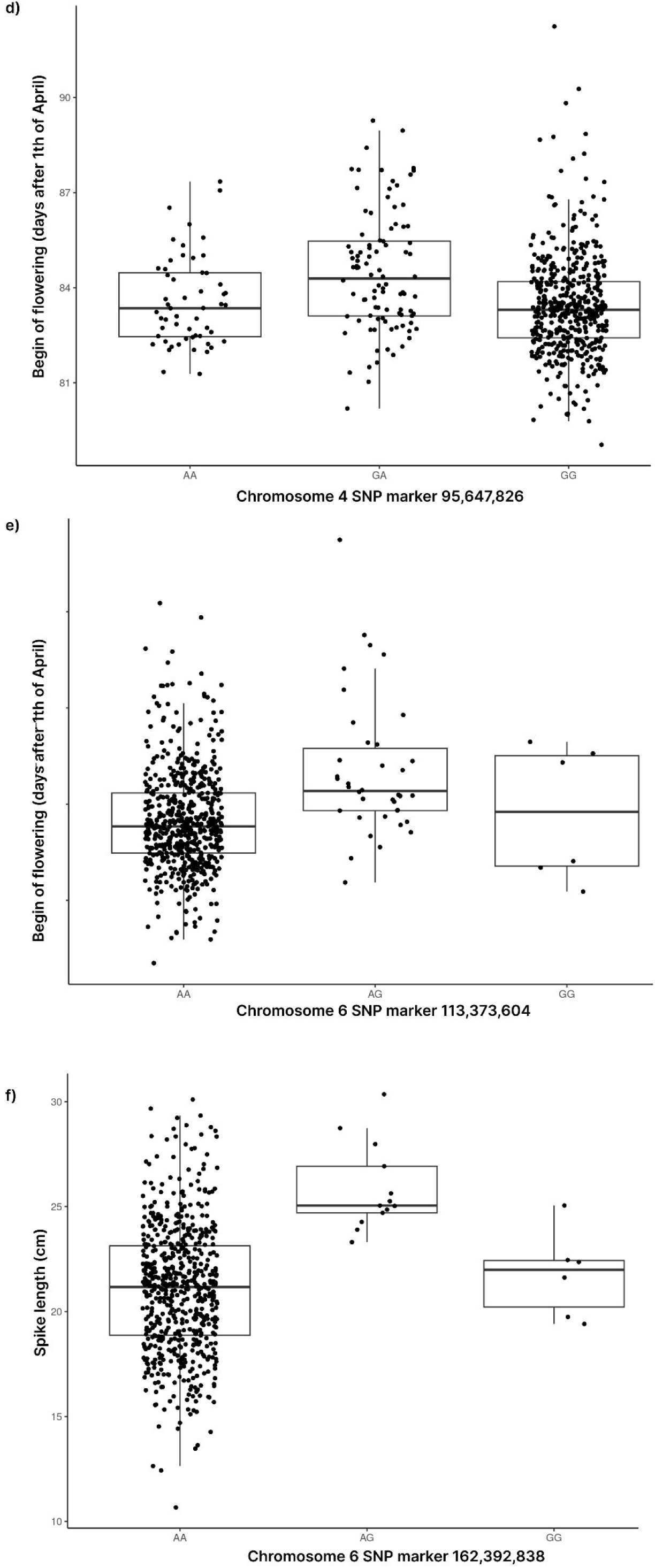
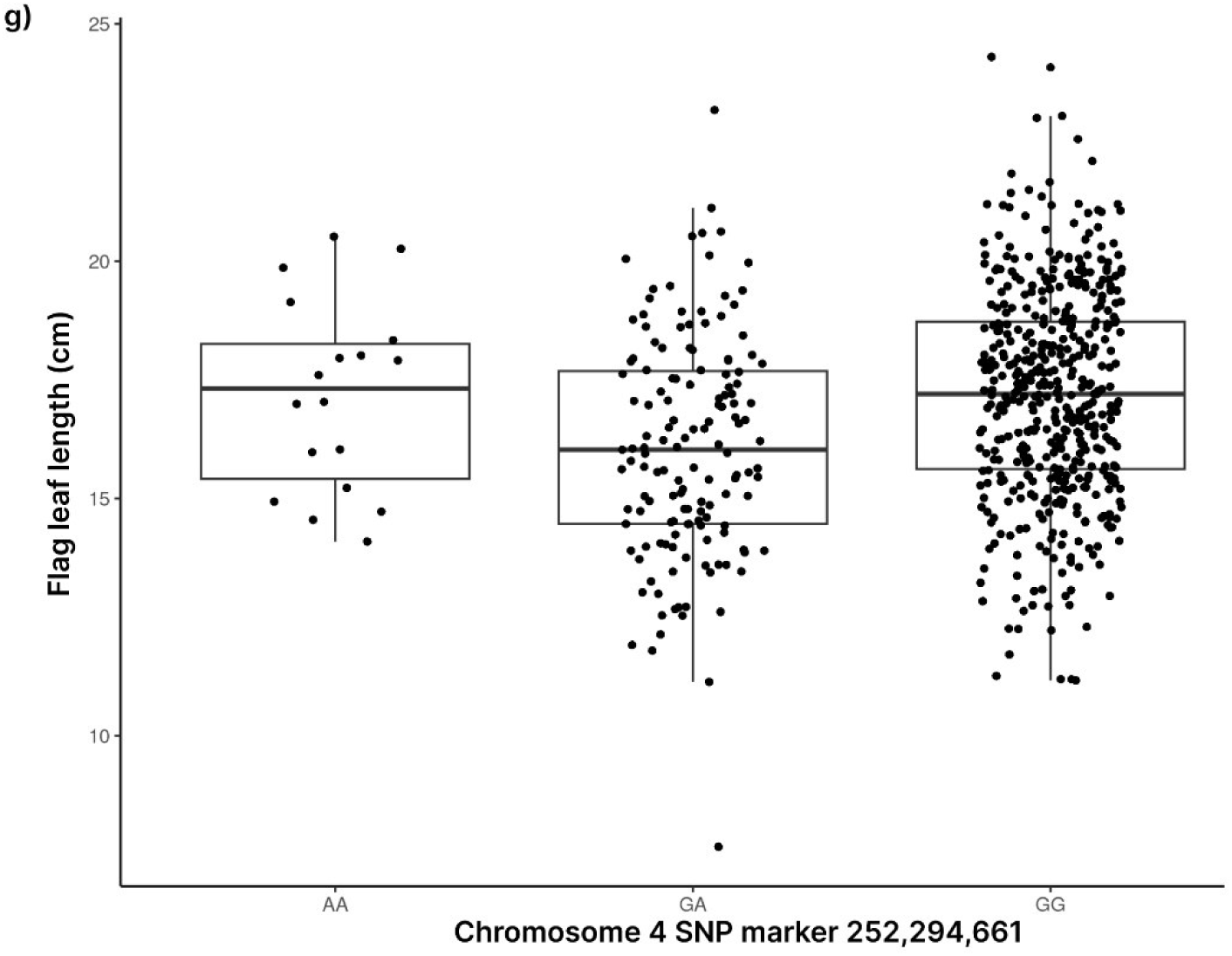
Effect plot for significantly associated markers for seed shattering a) on chromosome 7 on position 208,458,663, for seed yield b) on chromosome 1 on position 112,230,539, for flowering time c) on chromosome 4 on position 57,744,424, d) on chromosome 4 on position 95,647,826, e) on chromosome 6 on position 113,373,604, for spike length f) on chromosome 6 on position 162,392,838, and for flag leaf length g) on chromosome 4 on position 252,294,661. The x-axis shows the genotype for that marker and the y-axis the best-linear-unbiased estimators (BLUEs) for the respective traits over three environments phenotyping rust occurrence.

